# Repurposing of drugs as novel influenza inhibitors from clinical gene expression infection signatures

**DOI:** 10.1101/401315

**Authors:** Andrés Pizzorno, Olivier Terrier, Claire Nicolas de Lamballerie, Thomas Julien, Blandine Padey, Aurélien Traversier, Magali Roche, Marie-Eve Hamelin, Chantal Rhéaume, Séverine Croze, Vanessa Escuret, Julien Poissy, Bruno Lina, Catherine Legras-Lachuer, Julien Textoris, Guy Boivin, Manuel Rosa-Calatrava

**Affiliations:** Virologie et Pathologie Humaine - VirPath team, Centre International de Recherche en Infectiologie (CIRI), INSERM U1111, CNRS UMR5308, ENS Lyon, Université Claude Bernard Lyon 1, Université de Lyon, Lyon 69008, France; Research Center in Infectious Diseases of the CHU de Quebec and Laval University, Quebec City, QC G1V 4G2, Canada; Viroscan3D SAS, Lyon 69008, France; VirNext, Faculté de Médecine RTH Laennec, Université Claude Bernard Lyon 1, Université de Lyon, Lyon 69008, France; ProfileXpert, SFR-Est, CNRS UMR-S3453, INSERM US7, Université Claude Bernard Lyon 1, Université de Lyon, Lyon 69008, France; Laboratoire de Virologie, Centre National de Référence des virus Influenza Sud, Institut des Agents Infectieux, Groupement Hospitalier Nord, Hospices Civils de Lyon, Lyon F-21 France; Pôle de Réanimation, Hôpital Roger Salengro, Centre Hospitalier Régional 22 et Universitaire de Lille, Université de Lille 2, Lille 59000, France; Ecologie Microbienne, UMR CNRS 5557, USC INRA 1364, Université Claude Bernard Lyon 1, Université de Lyon, Villeurbanne 69100, France; Service d’Anesthésie et de Réanimation, Hôpital Edouard Herriot, Hospices Civils de Lyon, Lyon 69003, France; Pathophysiology of Injury-Induced Immunosuppression (PI3), EA 7426 Hospices Civils de Lyon,bioMérieux, Université Claude Bernard Lyon 1, Hôpital Edouard Herriot, Lyon 69003, France

**Keywords:** Influenza viruses, antivirals, inhibitors of viral infection, transcriptome, host targeting, drug repurposing

## Abstract

**Background:** Influenza virus infections remain a major and recurrent public health burden. The intrinsic ever-evolving nature of this virus, the suboptimal efficacy of current influenza inactivated vaccines, as well as the emergence of resistance against a limited antiviral arsenal, highlight the critical need for novel therapeutic approaches. In this context, the aim of this study was to develop and validate an innovative strategy for drug repurposing as host-targeted inhibitors of influenza viruses and the rapid evaluation of the most promising candidates in Phase II clinical trials.

**Methods:** We exploited *in vivo* global transcriptomic signatures of infection directly obtained from a patient cohort to determine a shortlist of already marketed drugs with newly identified, host-targeted inhibitory properties against influenza virus. The antiviral potential of selected repurposing candidates was further evaluated *in vitro, in vivo* and *ex vivo*.

**Results:** Our strategy allowed the selection of a shortlist of 35 high potential candidates out of a rationalized computational screening of 1,309 FDA-approved bioactive molecules, 31 of which were validated for their significant *in vitro* antiviral activity. Our *in vivo* and *ex vivo* results highlight diltiazem, a calcium channel blocker currently used in the treatment of hypertension, as a promising option for the treatment of influenza infections. Additionally, transcriptomic signature analysis further revealed the so far undescribed capacity of diltiazem to modulate the expression of specific genes related to the host antiviral response and cholesterol metabolism. Finally, combination treatment with diltiazem and virus-targeted oseltamivir neuraminidase inhibitor further increased antiviral efficacy, prompting rapid authorization for the initiation of a Phase II clinical trial.

**Conclusions:** This original, host-targeted, drug repurposing strategy constitutes an effective and highly reactive process for the rapid identification of novel anti-infectious drugs, with potential major implications for the management of antimicrobial resistance and the rapid response to future epidemic or pandemic (re)emerging diseases for which we are still disarmed.

## Background

Besides their well-known pandemic potential, annual outbreaks caused by influenza viruses account for several million respiratory infections and 250,000 to 500,000 deaths worldwide [1]. This global high morbidity and mortality of influenza infections represents a major and recurrent public health threat with high economic burden. In this context, the suboptimal vaccine coverage and efficacy, coupled with recurrent events of viral resistance against a very limited antiviral portfolio, emphasize an urgent need for innovative treatment strategies presenting fewer obstacles for their clinical use [2].

For decades, the strategy for antiviral development was mostly based on serial screenings of hundreds of thousands of molecules to identify “hits” and ‘leads” that target specific viral determinants, a quite costly and time-consuming process. However, the dramatic reduction in successful candidate identification over time [3], along with a concomitant increase of regulatory complexity to implement clinical trials, have fostered rising interest in novel strategies. Indeed, new approaches, focused on targeting the host instead of the virus, as well as on marketed drug repurposing for new antiviral indications [3–5] have been recently proposed in the context of global health emergencies posed by Ebola [6] and Zika [7] viruses. Such innovative strategies are strongly supported by a shift of paradigms in drug discovery, from “one-drug-one-target” to “one-drug-multiple-targets” [8]. In that sense, different *in silico* approaches based on structural bioinformatic studies [9, 10], systems biology approaches [11] and host gene expression analyses [12] have been applied to decipher multi-purpose effects of many US Food and Drug Administration (FDA)-approved drugs. Additionally, as successfully demonstrated in antiretroviral therapy [13], targeting host instead of viral determinants may confer a broad-spectrum antiviral efficacy, and also reduce the risk of emergence of drug resistance against influenza viruses [14]. As a result, the last decade has witnessed several host-directed experimental approaches against influenza infections, notably nitazoxanide, DAS181 or acetylsalicylic acid [15–17].

In line with this emerging trend, we previously postulated that host global gene expression profiling can be considered as a “fingerprint” or signature of any specific cell state, including during infection or drug treatment, and hypothesized that the screening of databases for compounds that counteract virogenomic signatures could enable rapid identification of effective antivirals [18]. Based on this previous proof-of-concept obtained from *in vitro* gene expression profiles, we further improved our strategy by analyzing paired upper respiratory tract clinical samples collected during the acute infection and after recovery from a cohort of influenza A(H1N1)pdm09-infected patients and determined their respective transcriptomic signatures. We then performed an *in silico* drug screening using Connectivity Map (CMAP), the Broad Institute’s publicly available database of more than 7,000 drug-associated gene expression profiles [19, 20], and identified a list of candidate bioactive molecules with signatures anti-correlated with those of the patient’s acute infection state (**Figure 1A**). The potential antiviral properties of selected FDA-approved molecules were firstly validated *in vitro*, and the most effective compounds were further compared to oseltamivir for the treatment of influenza A(H1N1)pdm09 virus infections in both C57BL/6 mice and 3D reconstituted human airway epithelia. Altogether, our results highlight diltiazem, a calcium channel blocker with so far undescribed capacity to stimulate the epithelial antiviral defense, as a promising repurposed host-targeted inhibitor of influenza infection. Moreover, our results plead in favor of the combination of diltiazem with the virus-targeted antiviral oseltamivir for the improvement of current anti-influenza therapy, and possibly decreasing the risk of antiviral resistance. This study confirms the feasibility and interest of integrating clinical virogenomic and chemogenomic inputs as part of a drug repurposing strategy to accelerate bedside-to-bench and bench-to-bedside drug development.

**Fig. 1.**
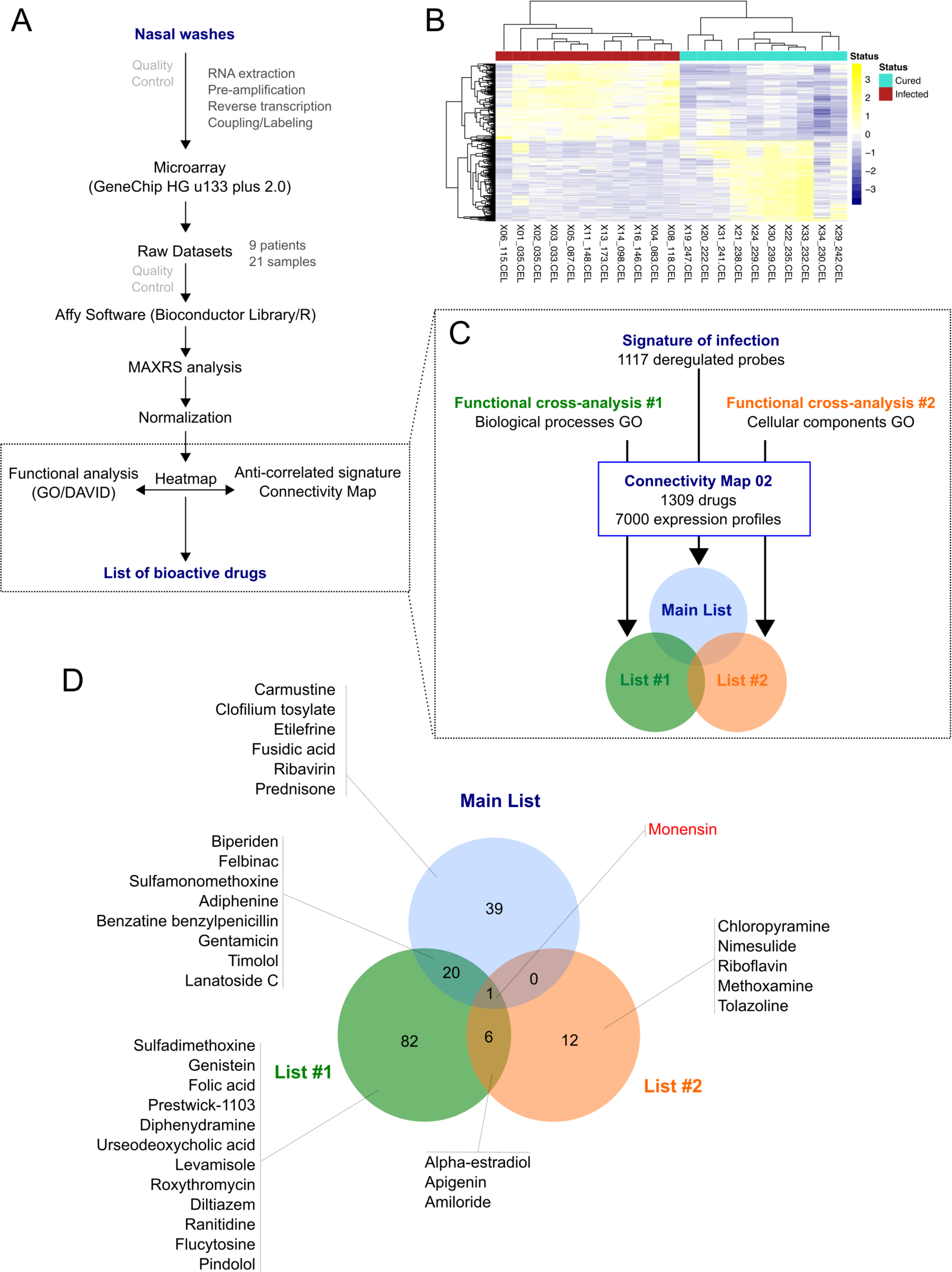
From nasal wash clinical samples to a shortlist of 35 candidate molecules. **(A)** Overview of the *in silico* strategy used in this study. A detailed description of the strategy is described in the Online Methods section. **(B)** Hierarchical clustering and heatmap of the 1,117 most differentially deregulated genes between “infected” (red) and “cured” (light green) samples. Raw median centered expression levels are color coded from blue to yellow. Dendrograms indicate the correlation between clinical samples (columns) or genes (rows). **(C)** Functional cross-analysis of candidate molecules obtained from Connectivity Map (CMAP). Three lists of candidate molecules were obtained using different set of genes in order to introduce functional bias and add more biological significance to this first screening: a Main List based on the complete list of differentially expressed genes, and two other lists (List #1 and #2) based on subsets of genes belonging to significantly enriched Gene Ontology (GO) terms. **(D)** Venn Diagram comparing the total 160 molecules obtained from the three lists described in (C), with monensin as the only common molecule. Only the candidates selected for *in vitro* screening and validation are depicted.

## Methods

### Clinical samples

A previously published randomized clinical trial (ClinicalTrials.gov identifier NCT00830323) was conducted in Lyon and Paris (France) during the peak circulation of the influenza A(H1N1)pdm09 virus, with the aim to assess the efficacy of oseltamivir-zanamivir combination therapy compared with oseltamivir monotherapy [21]. Briefly, patients tested positive for influenza A infection by the QuickVue rapid antigen kit (Quidel) were randomized in one of the two treatment groups and nasal wash specimens were collected within two hours of the first visit and every 24 h until 96 h after treatment initiation. Nasal swabs were also performed on days 5 and 7. In voluntary patients, an optional supplementary nasal wash was performed at least three months after influenza infection (recovery phase). H1N1 subtype was further confirmed by PCR. For nine of these patients, transcriptomic data were obtained from paired samples collected during influenza infection without treatment and in the recovery phase.

### Sample processing, RNA preparation and hybridization

Nasal wash samples were collected in RNAlater^®^ Stabilization Solution (Thermo Fisher Scientific). Total RNA was extracted using RNeasy Micro kit (Qiagen) following the manufacturer’s instructions. RNA quality was assessed using a Bioanalyzer2100 (Agilent technologies, Inc, Palo Alto, CA, USA). To account for samples having low amount and/or partially degraded RNA (RNA Integrity Numbers between 1 and 8), we applied two types of corrections: i) cRNA labelling was performed after a linear amplification protocol, as previously described [22] and ii) raw signals obtained after hybridization of labelled cRNA on microarray and data acquisition were processed using the MAXRS algorithm [23]. Labeled cRNA were hybridized on Affymetrix HG-U133plus2 microarrays according to manufacturer’s instructions in a GeneChip^®^ Hybridization Oven 640 (Affymetrix) and microarrays were subsequently scanned in an Affymetrix 3000 7G scanner.

### Data normalization & MAXRS computational analysis

The MAXRS algorithm [23] is particularly suited to gene expression analysis under low hybridization conditions. Briefly, this method takes advantage of the specific design of Affymetrix probe sets, which are composed of an average of 11 different probes that target the same locus, and is based on the observation that for most of the probe sets the same probe shows the highest fluorescence intensity in almost all arrays. For each microarray (m = 1..M) and for each probeset (t = 1..T), fluorescence intensity values on microarray m of all probes (p = 1..P_t_) belonging to the probeset t are sorted in increasing order. These ranks are denoted as r_mtp_. Then, we calculated across all microarrays the rank sum (RS_tp_) for each probeset t for each probe p belonging to the probeset t. Finally, for each probeset t, we kept the three probes p with the highest RS_tp_. The mean intensity of these three probes is attributed to the probeset t. As it is common practice with many modern pre-processing algorithms, and because of the low global fluorescence signal intensity, mismatched probes were excluded from MAXRS analysis.

After pre-processing the raw dataset with the MAXRS algorithm, a normalization step was performed using Tukey median-polish algorithm [24]. Differential expression was assessed by applying a Student t-test for each probeset, and multiple testing was corrected using the Benjamini-Hochberg algorithm in the qvalue library [25]. For further downstream analysis, genes were selected according to two criteria: i) absolute fold change >2, and ii) corrected p-value <0.05. Data were generated according to the Minimum Information About a Microarray Experiment guidelines and deposited in the National Center for Biotechnology Information’s Gene Expression Omnibus [26] under accession number GSE93731.

### Functional analysis

Functional enrichment analysis was performed on a selection of differentially-expressed genes with DAVID tools [27], using the Gene Ontology (GO) [28]. To further select genes for the CMAP query, we selected 6 Biological Process terms (GO_BP: GO:0009615-response to virus; GO:0006955-immune response; GO:0042981-regulation of apoptosis; GO:0006952-defense response; GO:0009611-response to wounding; GO:0042127-regulation of cell proliferation) that shared > 90% of genes with all significantly enriched GO_BP terms, and 3 relevant Cellular Component terms (GO_CC: GO:0031225-anchored to membrane; GO:0005829-cytosol; GO:0005654-nucleoplasm). To visualize and compare the different lists of compounds, Venn diagrams were obtained using the webtool developed by Dr. Van de Peer’s Lab at Ghent University (http://bioinformatics.psb.ugent.be/webtools/Venn/).

### Cells and viruses

Human lung epithelial A549 cells (ATCC CCL-185) were maintained in Dulbecco’s modified Eagle’s medium (DMEM) supplemented with 10 % fœtal calf serum and supplemented with 2 mM L-glutamine (Sigma Aldrich), penicillin (100 U/mL) and streptomycin (100 µg/mL) (Lonza), maintained at 37 °C and 5% CO_2_.

Influenza viruses A/Lyon/969/09 and A/Quebec/144147/09 were produced in MDCK (ATCC CCL-34) cells in EMEM supplemented with 2 mM L-glutamine (Sigma Aldrich), penicillin (100 U/mL), streptomycin (100 µg/mL) (Lonza) and 1 µg/mL trypsin. Viral titers in plaque forming units (PFU/ml) and tissue culture infectious dose 50% (TCID50/mL) were determined in MDCK cells as previously described [29, 30].

### Viral growth assays

For viral growth assays in the presence of molecules, A549 cells were seeded 24 h in advance in multi-well 6 plates at 1.8 x 10^5^ cells/well. Three treatment protocols were evaluated. 1) In pre-treatment protocol, cells were washed with DMEM and then incubated with different concentrations of candidate molecules diluted in DMEM supplemented with 2 mM L-glutamine (Sigma Aldrich), penicillin (100 U/mL), streptomycin (100 µg/mL) (Lonza) and 0.5 µg/mL trypsin. Six hours after treatment, cells were washed and then infected with A/Lyon/969/09 (H1N1)pdm09 virus at a MOI of 0.1. 2) In pre-treatment plus post-treatment protocol, cells were initially treated and infected in the same conditions as explained above. One hour after viral infection, a second identical dose of candidate molecules in supplemented DMEM was added. 3) In post-treatment protocol, cells without pre-treatment were infected in the conditions described and treatments with candidate molecules at the indicated concentrations were initiated 24 h p.i. In all cases, supernatants were collected at 48 h p.i. and stored at -80 °C for TCID50/ml viral titration.

### Viability and cytotoxicity assays

Cell viability was measured using the CellTiter 96^®^ AQueous One Solution Cell Proliferation Assay (MTS, Promega). A549 cells were seeded into 96-well plates and treated with different concentrations of molecules or solvents. Cells were incubated at 37 °C and 5% CO_2_ and then harvested at different time-points, following the same scheme as in viral growth assays. Results were presented as a ratio of control values obtained with solvents. Treatment-related toxicity in human airway epithelia (HAE) was measured using the Cytotoxicity Detection Kit^PLUS^ (LDH, Roche) according to the manufacturer’s instructions. Briefly, duplicate 100 µL-aliquots of basolateral medium from treated and control HAEs were incubated in the dark (room temperature, 30 minutes) with 100 µL of lactate dehydrogenase (LDH) reagent in 96-well plates. After incubation, “stop solution” was added and the absorbance was measured in a conventional microplate ELISA reader. The photometer was set up for dual readings to determine non-specific background at 750 nm, and absorbance was measured at 490 nm. Percent cytotoxicity was calculated as indicated by the manufacturer, using mock-treated and 1% triton-treated epithelia as “low” and “high” controls, respectively. Percent viability is presented as 100 – percent cytotoxicity.

### Mouse model of viral infection

All protocols were carried out in seven to nine-week old female C57BL/6N mice (Charles River, QC, Canada). Animals were randomized in groups of 15 according to their weight to ensure comparable median values on each group, and then housed in micro-isolator cages (5 animals per cage) in a biosafety 2 controlled environment (22 °C, 40% humidity, 12:12 h photoperiods), with *ad libitum* access to food and water.

On day 0, mice were lightly anesthetized with inhaled 3% isoflurane/oxygen, and then infected by intranasal (i.n.) instillation of influenza A/Quebec/144147/09 (H1N1)pdm09 virus in 30 µl of saline, as specified in each case. Control animals were mock-infected with 30 µl of saline. Candidate molecules were evaluated in two different treatment protocols: i) treatments were started on the same day of infection (day 0, 6 h prior to infection), or ii) treatments were started 24 h after infection (day 1). Regardless of treatment initiation time, all treatments were performed *per os* (150-µl gavage) once daily for 5 consecutive days (5 drug administrations in total). Mortality, body weight and clinical signs such as lethargy and ruffled fur were daily monitored on 10 animals/group for a total of 14 days. Animals were euthanized if they reached the humane endpoint of >20% weight loss. The remaining 5 animals/group were euthanized on day 5 p.i. to measure LVTs.

Vehicle (saline) or oseltamivir were used as placebo and positive treatment control, respectively. The oseltamivir dose (10 mg/kg/day) was adjusted to confer ∼50% protection in the selected experimental conditions and is considered a good correlate of half the normal dose of 150 mg/day given to humans [31]. The doses of repurposed candidate molecules were selected to be in the non-toxic range for mouse studies, according to published preclinical data for their first therapeutic indication. To validate this choice in our specific model, potential drug toxicity was evaluated in mock-infected animals treated with the same regimens as virus-infected mice.

### Pulmonary viral titers

In order to evaluate the effect of different treatments on viral replication, 5 animals per group were euthanized on day 5 p.i. and lungs were removed aseptically. Mice were randomly selected from the 3 cages of each group to minimize cage-related bias. Lungs were homogenized in 1 ml of PBS using a bead mill homogenizer (Tissue Lyser, Qiagen) and debris was pelleted by centrifugation (2,000 g, 5 min). Triplicate 10-fold serial dilutions of each supernatant were plated on ST6GalIMDCK cells (kindly provided by Dr. Y. Kawaoka, University of Wisconsin, Madison, WI) and titrated by plaque assays [29]. The investigator was blinded to group allocation.

### Viral infection in reconstituted human airway epithelium (HAE)

MucilAir^®^ HAE were obtained from Epithelix SARL (Geneva, Switzerland) and maintained in air-liquid interphase with specific culture medium in Costar Transwell inserts (Corning, NY, USA) according to manufacturer’s instructions. For infection experiments, apical poles were gently washed with warm PBS and then infected with a 100-µL dilution of influenza A/Lyon/969/09 (H1N1)pdm09 virus in OptiMEM medium (Gibco, ThermoFisher Scientific) at a MOI of 0.1. Basolateral pole sampling as well as 150-µL OptiMEM apical washes were performed at the indicated time points, and then stored at -80 °C for PFU/mL and TCID50/mL viral titration. Treatments with specific dilutions of candidate molecules alone or combined with oseltamivir in MucilAir^®^ culture medium were applied through basolateral poles. All treatments were initiated on day 0 (5 h after viral infection) and continued once daily for 5 consecutive days (5 drug administrations in total). Variations in transepithelial electrical resistance (Δ TEER) were measured using a dedicated volt-ohm meter (EVOM2, Epithelial Volt/Ohm Meter for TEER) and expressed as Ohm/cm^2^.

### High throughput sequencing and bioinformatics analysis

cDNA libraries were prepared from 200 ng of total RNA using the Scriptseq™ complete Gold kit-Low Input (SCL6EP, Epicentre), according to manufacturer’s instructions. Each cDNA library was amplified and indexed with primers provided in the ScriptSeq™ Index PCR Primers kit (RSBC10948, Epicentre) and then sequenced as 100 bp paired-end reads. Prior to sequencing, libraries were quantified with QuBit and Bioanalyzer2100, and indexed libraries were pooled in equimolar concentrations. Sequencing was performed on an Illumina HiSeq 2500 system (Illumina, Carlsbad, CA), with a required minimum of 40 million reads sequenced per sample. Conversion and demultiplexing of reads was performed using bcl2fastq 1.8.4 (Illumina). The FastQC software (http://www.bioinformatics.babraham.ac.uk/projects/fastqc) was used for quality controls of the raw data. Reads were trimmed using the Trimmomatic [32] software, with a minimum quality threshold of Q30. Trimmed reads were pseudo-aligned to the *Homo sapiens* genome (GRCh38.p11) using the Kallisto software [33]. Statistical analysis was performed in R3.3.1 with the package EdgeR 3.14.0 [34]. Differential expression was calculated by comparing each condition to the mock using a linear model. The Benjamini-Hochberg procedure was used to control the false discovery rate (FDR). Transcripts with an absolute fold change >2 and a corrected p-value <0.05 were considered to be differentially expressed. Enriched pathways and GO terms were assessed with DAVID 6.8 [27]. For visualization purposes, a heatmap and stacked barplots were constructed in R3.3.1 on mean-weighted fold changes and association between conditions were assessed by Spearman correlation analysis.

### Statistical analysis

All experimental assays were performed in duplicate at a minimum, and representative results are shown unless indicated otherwise. No statistical methods were used to predetermine sample size in animal studies, which were estimated according to previous studies and the known variability of the assays. No mice were excluded from post-protocol analyses, the experimental unit was an individual animal and equal variance was assumed. Kaplan-Meier survival plots were compared by Log-Rank (Mantel-Cox) test and hazard ratios (HR) were computed by the Mantel-Haenszel method. Weight loss and viral titers of all groups were compared by one-way analysis of variance (ANOVA) with Tukey’s multiple comparison post-test. The testing level (α) was 0.05. Statistical analyses were performed on all available data, using GraphPad, Prism 7.

## Results

### Generation of clinical virogenomic profiles

We determined *in vivo* transcriptional signatures of infection from paired nasal wash samples of nine untreated patients, collected during acute A(H1N1)pdm09 pandemic influenza infection (“infected”) and at least three months later to ensure a recovery non-infected state (“cured”) [21]. The nine patients from whom transcriptomic data could be obtained constitute a representative sample of the whole studied cohort, except for the male sex ratio (**Table S1**). We combined two strategies to tackle the characteristic low RNA amount/quality of this type of clinical samples. Firstly, cRNA labelling was performed after a linear amplification of initial RNA, as previously described [35]. Secondly, raw signals obtained after hybridization of labelled cRNA on microarray and data acquisition were processed using the MAXRS algorithm [23] to overcome low hybridization conditions. This approach, initially developed for the analysis of heterologous hybridizations, takes advantage of the specific design of the Affymetrix^®^ microarray used in our study, with several probes targeting the same locus [23].

After normalization, differentially expressed genes were selected based on two criteria: i) an absolute fold change >2, and ii) a Benjamini-Hochberg corrected p-value <0.05. We therefore identified a total of 1,117 commonly deregulated probes, with almost equal proportion of up-regulated (48.4%; n=541) and down-regulated probes (51.7%; n=576). Remarkably, despite considerable inter-patient variability among recovery state samples, a substantial homogenization of transcriptional profiles was observed in the context of infection, as shown in the heatmap presented in **Figure 1B** and by the median Spearman’s ρ correlation values for both groups (0.60 “cured” vs. 0.90 “infected”). These virogenomic signatures of infection constituted the input for the subsequent *in silico* query for the identification of candidate compounds.

### *In silico* cross-analysis of chemogenomic versus virogenomic clinical profiles

We then performed an *in-silico* search for molecules that reverse the virogenomic signature of infection, using the CMAP database (Build 02) as previously described [18]. CMAP is a collection of genome-wide transcriptional expression data from cultured human cells treated with bioactive small molecules. HG-U133plus2 probesets were mapped to the U133A probesets using the Ensembl BioMarts online tool [36, 37], and connectivity scores and p-values were obtained using the CMAP algorithm [19, 20]. With the global set of 1,000 most differentially expressed genes as input (**Figure 1C**, Main List), we obtained a preliminary list of 60 candidate compounds. In parallel, we used two other subsets of genes belonging to significantly enriched Gene Ontology (GO) terms obtained from microarray analyses to introduce functional bias and add more biological significance to our first screening. Hence, by using 6 Biological Process terms (GO_BP) that shared more than 90% of genes (**Figure 1C**, Functional cross-analysis #1), a second list of 109 compound candidates was obtained. A third list of 19 compounds was obtained using 3 relevant Cellular Component terms (GO_CC) (**Figure 1C**, Functional cross-analysis #2). The comparison of the 160 compounds from the three distinct lists (12.2% of compounds of CMAP, **Table S2**) highlighted monensin as the only common compound (**Figure 1D**).

To rationally reduce the number of drug candidates, bioactive drugs were excluded if not compatible with a final use as antiviral, mostly for safety (e.g. teratogens, intercalating agents) and/or pharmacological (e.g. documented low bioavailability) reasons, based on clinical data and the PubMed/PubChem databases. Thus, the number of candidates was initially decreased to 139 and then to 110 (**Figure S1**). We subsequently determined a shortlist of 35 bioactive molecules (<3% of CMAP, **Table 1**) for *in vitro* screening, based on two main criteria: i) molecules representative of the different pharmacological classes identified, and ii) molecules evenly distributed in the three lists obtained after *in silico* screening (Main List, List #1 and List #2, **Figure 1D**), which comprise a panoply of documented pharmacological classes, including anti-fungal agents (e.g. monensin, flucytosine), anti-inflammatory agents (e.g. felbinac, apigenin, prednisone) and adrenergic agonists/antagonists (timolol, methoxamine, tolazoline), as represented in the Venn diagram (**Figure 1D**, **Table 1**). Interestingly, at least 14 (40%) molecules from our short-list belong to a pharmacological class related with anti-microbial or anti-inflammatory activities (**Table 1**, #), and 11 (31.4%) have already been reported in the literature for their antiviral properties against influenza or other viruses (**Table 1**, *), notably the nucleoside inhibitor ribavirin [38, 39] and the ionophore monensin [40].

**Table 1.** Shortlist of the 35 selected molecules and their documented pharmacological classes. Shortlist of the 35 selected candidates representative of the 110 molecules obtained from the *in silico* screening (Fig 1 and S1). Documented pharmacological classes were obtained from PubChem (https://pubchem.ncbi.nlm.nih.gov). Asterisks (*) indicate molecules previously evaluated for their antiviral properties according to the literature, and numerals (#) those belonging to anti-microbial or anti-inflammatory related pharmacological classes

| Name | Pharmacological Class |
| --- | --- |
| Adiphenine | Parasympatholytics/Anticholinergics/Antispasmodics |
| Alpha-estradiol* | 5 alpha-reductase inhibitors/Androgenic alopecia treatment |
| Amiloride*# | Epithelial Sodium Channel Blockers/Diuretics/Acid Sensing Ion Channel Blockers |
| Apigenin*# | Anti-Inflammatory Agents, Non-Steroidal/? |
| Benzathine benzylpenicillin# | Anti-Bacterial Agents |
| Biperiden* | Antiparkinson Agents/Muscarinic Antagonists/Parasympatholytics |
| Carmustine | Antineoplastic Agents, Alkylating |
| Chloropyramine | Histamine H1 Antagonists |
| Clofilium tosylate | Anti-Arrhythmia Agents |
| Diltiazem | Antihypertensive Agents/Calcium Channel Blockers/Cardiovascular Agents/Vasodilator Agents |
| Diphenhydramine | Anesthetics, Local/Anti-Allergic Agents/Antiemetics/Histamine H1 Antagonists/Hypnotics and Sedatives |
| Etilefrine | Adrenergic beta-1 and alpha agonist /Cardiotonic/antihypotensive agent. |
| Felbinac# | Anti-Inflammatory Agents, Non-Steroidal |
| Flucytosine*# | Antifungal Agents/Antimetabolites |
| Folic acid | Hematinics/Vitamin B Complex |
| Fusidic acid# | Anti-Bacterial Agents/Protein Synthesis Inhibitors |
| Genistein | Anticarcinogenic Agents/Phytoestrogens/Protein Kinase Inhibitors |
| Gentamicin# | Anti-Bacterial Agents/Protein Synthesis Inhibitors |
| Lanatoside C | Anti-Arrhythmia Agents |
| Levamisole* | Adjuvants, Immunologic/Antinematodal Agents/Antirheumatic Agents |
| Methoxamine | Adrenergic alpha-1 Receptor Agonists/Sympathomimetics/Vasoconstrictor Agents |
| Monensin*# | Antifungal Agents/Antiprotozoal Agents/Coccidiostats/Proton Ionophores/Sodium Ionophores |
| Nimesulide*# | Anti-Inflammatory Agents, Non-Steroidal/Cyclooxygenase Inhibitors |
| Pindolol | Adrenergic beta-Antagonists/Antihypertensive Agents/Serotonin Antagonists/Vasodilator Agents |
| Prednisone# | Anti-Inflammatory Agents/Antineoplastic Agents, Hormonal/Glucocorticoids |
| Prestwick-1103# | Anti-Inflammatory Agents, Non-Steroidal/Cyclooxygenase Inhibitors |
| Ranitidine | Anti-Ulcer Agents/Histamine H2 Antagonists |
| Ribavirin* | Antimetabolites/Antiviral Agents |
| Riboflavin* | Photosensitizing Agents/Vitamin B Complex |
| Roxithromycin# | Anti-Bacterial Agents |
| Sulfadimethoxine# | Anti-Infective Agents |
| Sulfamonomethoxine# | Anti-Infective Agents |
| Timolol* | Adrenergic beta-Antagonists/Anti-Arrhythmia Agents/Antihypertensive Agents |
| Tolazoline | Adrenergic alpha-Antagonists/Antihypertensive Agents/Vasodilator Agents |
| Ursodeoxycholic acid | Cholagogues and Cholaretics |

### Inhibitory effect of the selected molecules on A(H1N1)pdm09 viral growth *in vitro*

*In vitro* screening of the antiviral potency of the 35 selected molecules was performed in A549 human lung epithelial cells seeded in 6-well plates. Firstly, we evaluated the impact of 6 h pre-treatment with a 10-fold drug concentration range, using the original CMAP concentration as reference. Six hours after treatment, cells were washed and infected with influenza A(H1N1)pdm09 virus at a multiplicity of infection (MOI) of 0.1. Viral titers in supernatants collected from treated samples at 48 h post infection (p.i.) were normalized with those measured in mock-treated controls (>10^5^ TCID50/mL). Potential treatment-induced cell toxicity was evaluated in the same experimental conditions using the MTS assay and expressed also as the percentage of cell viability compared to non-infected controls (**Figure 2**). Based on antiviral activity and cell viability profiles obtained (**Figure 2A**, blue triangles), we defined as “inhibitors” compounds that fulfilled the following two criteria: i) induce >75% reduction on viral production, and ii) have minor impact on cell viability, with relative values in the 90%-110% range (**Figure 2A**, squares in **left panels** and zooms in **right panels**). A total of 10 compounds (28.6%) matched both criteria, mainly when used at a 10-fold CMAP concentration (**Figure 2B**), yet only a limited number of them exhibited classic dose-dependent inhibition. Whenever possible, as in the case of monensin or ranitidine for example, EC50 values were calculated, which were mostly in the micromolar range (**Figure 2B**).

In a second round of screening, we tested the same 6 h pre-treatment but with serial 10-fold dilutions from the initial CMAP concentration to CMAP/10,000, followed by one additional treatment immediately after infection (**Figure 2A**, green circles in **left** and **right panels**). In these conditions, 30 compounds (85.7%) met our criteria to be considered as inhibitors of viral production (**Figure 2C**), with half of them showing a classic dose-dependent inhibition effect. Calculated EC50 values were in the nanomolar range and hence significantly lower than those calculated in the context of pre-treatment only. Dose response curves and calculated EC50 for all the 35 compounds are presented in **Figure S2** and **Table S3**, respectively.

**Fig. 2.**
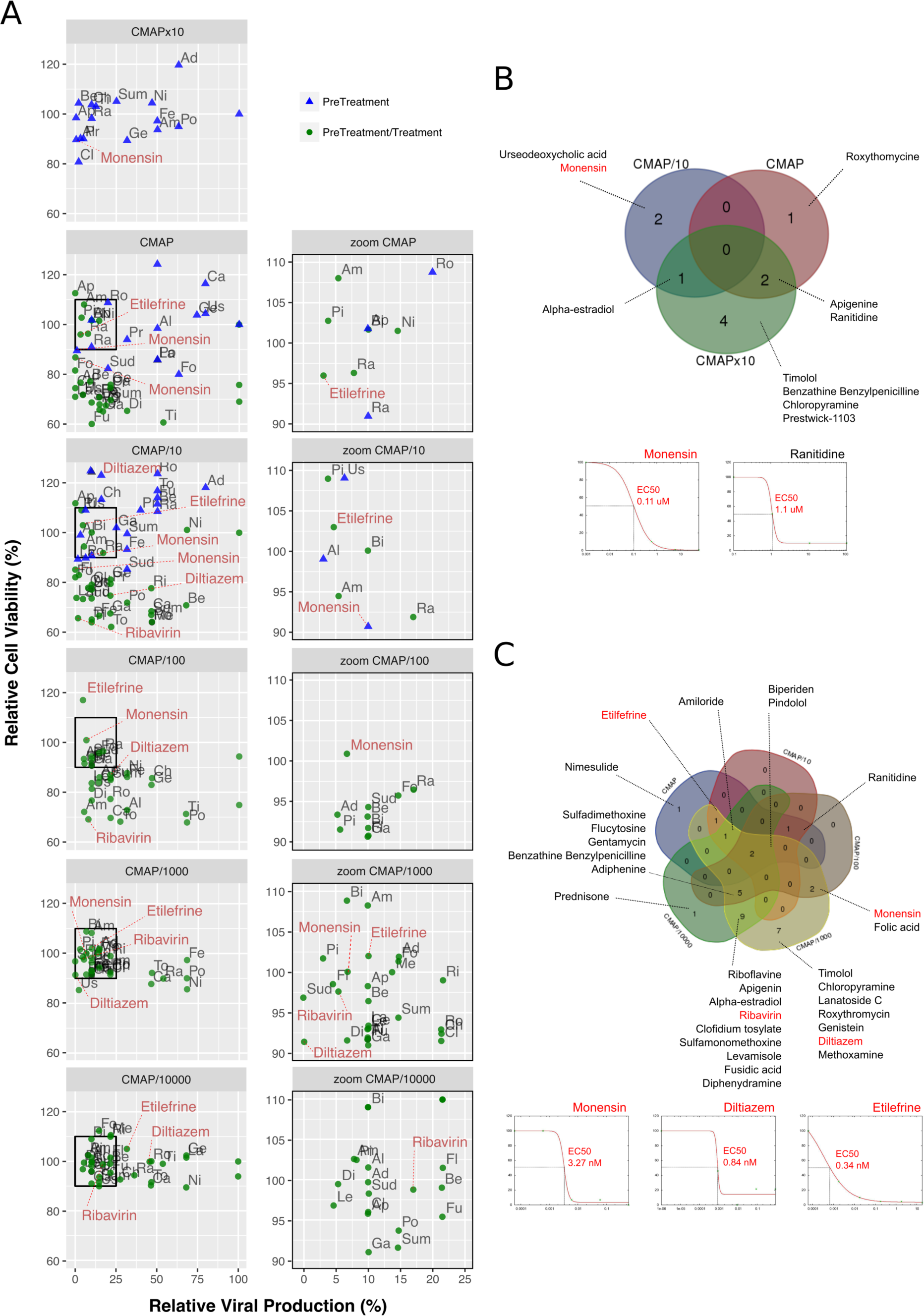
Screening and validation of the effect of selected molecules on A(H1N1)pdm09 viral growth *in vitro*. **(A, left)** Evaluation on A549 cells of the antiviral potency of the 35 candidates selected by *in silico* analysis. Relative viral production (%, X axis) and relative cell viability (%, Y axis) of both pre-treatment (blue triangles) and pre-treatment/treatment (green circles) regimens were evaluated. A 10-fold drug concentration range using CMAP as reference (CMAP x10, CMAP, CMAP/10, CMAP/100, CMAP/1,000 and CMAP/10,000) was used. CMAPx10 was only tested in the context of pre-treatment, by anticipation of a lower efficacy of molecules in this experimental setup. All experimental assays were performed in triplicate and mean values are represented. **(A, right)** Zoom panels depicting molecules defined as “inhibitors” according to the following two criteria: i) induce a 75% or higher reduction on viral production, and ii) have minor impact on cell viability, with relative values in the 90%-110% range. For clarity purposes, with the exception of diltiazem, etilefrine, monensin and ribavirin, abbreviations were used: Adiphenine "Ad”; Alpha-estradiol "Al"; Amiloride "Am"; Apigenin "Ap"; Benzathine Benzylpenicilline "Be"; Biperiden "Bi"; Carmustine "Ca"; Chloropyramine "Ch"; Clofidium tosylate "Cl"; Diphenydramine "Di"; Felbinac "Fe"; Flucytosine "Fl"; Folic acid "Fo"; Fusidic acid "Fu"; Genistein "Ge"; Gentamycin "Ga"; Lanatoside C "La"; Levamisole "Le"; Methoxamine "Me"; Nimesulide "Ni"; Pindolol "Pi"; Prednisone "Po"; Prestwick-1103 "Pr"; Ranitidine "Ra"; Riboflavine "Ri"; Roxythromycin "Ro"; Sulfadimethoxine "Sud"; Sulfamonomethoxine "Sum"; Timolol "Ti"; Tolazoline "To"; Urseodeoxycholic acid "Us". Dose-response curves for all the 35 molecules are presented in Supplementary Table 3. **(B)** Venn diagram of the 10 molecules identified in pre-treatment (10/35; 28.57%) and matching the “inhibitor” criteria, mainly when used at 10-fold CMAP concentration. EC50 curves for monensin and ranitidine are represented. **(C)** Venn diagram of the 30 “inhibitor” molecules identified in pre-treatment/treatment (30/35; 85.7%). EC50 curves for monensin, diltiazem and etilefrine are represented.

### Efficacy of selected molecules for the treatment of influenza A(H1N1)pdm09 virus infection in mice

Based on EC50 and cytoxicity data from the *in vitro* screening, we selected 8 molecules to investigate their potential as inhibitors of influenza A(H1N1)pdm09 in C57BL/6 mice. Oseltamivir, the standard antiviral for the treatment of influenza infections was used as control. All treatments were performed *per os*, starting 6 h before infection and being continued once daily for 5 consecutive days (5 drug administrations in total) (**Figure 3**). While animals treated with oseltamivir or monensin showed clinical improvement compared to the saline (placebo) group in terms of survival and weight loss (oseltamivir only), treatment with Lanatoside C, prednisolone, flucytosine, felbinac and timolol showed no clinical benefit at the selected concentrations (**Figure S5A**). In contrast, diltiazem and etilefrine not only significantly improved survival and maximum mean weight losses (**Figure 3A**-**B**), but also showed at least 1-log reductions in lung viral titers (LVTs) on day 5 p.i. (**Figure 3C**). Importantly, no signs of toxicity were observed for any of the drugs at the regimens tested (**Figure S3B**).

**Fig. 3.**
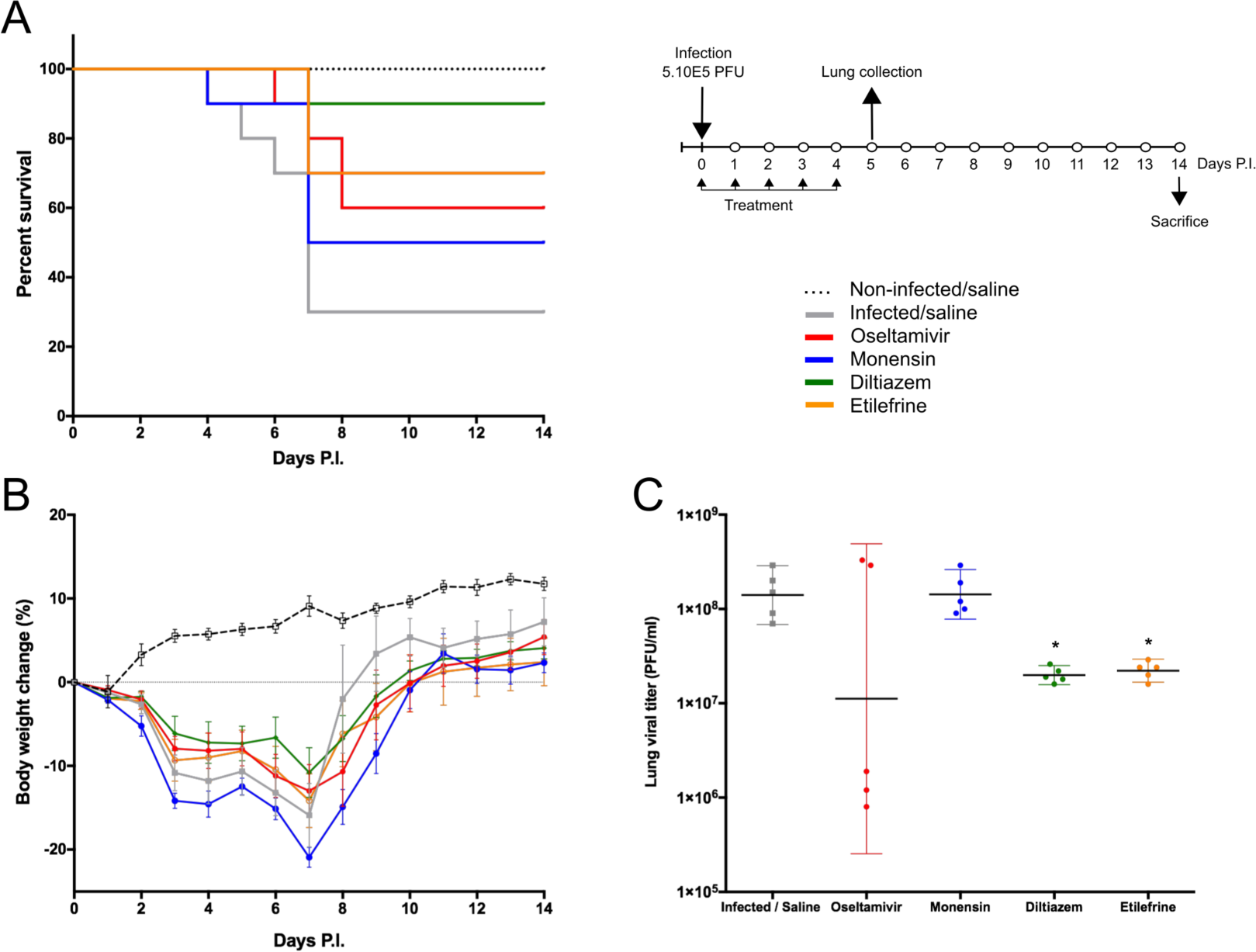
Efficacy of oral administration of selected molecules in mice infected with influenza A(H1N1)pdm09 virus. C57BL/6N mice (n=15/group) were intranasally inoculated with 5 x 10^5^ PFU of influenza A/Quebec/144147/09 virus on day 0 and treated by gavage with saline (grey), oseltamivir 10 mg/kg/day (red), monensin 10 mg/kg/day (blue), diltiazem 90 mg/kg/day (green), or etilefrine 3 mg/kg/day (orange). A mock-infected, saline-treated group (black dotted line, n=6) was included as control. Treatments were initiated on day 0 (6 h before infection) and administered once daily for 5 consecutive days. **(A)** Survival rates (n=10/group), **(B)** mean weight changes (±SEM, n=10/group or remaining mice) and **(C)** median (±CI95, n=5/group) lung viral titers on day 5 p.i. are shown. *p<0.05, **p<0.01 and ***p<0.001 compared to the infected saline-treated group by one-way ANOVA with Tukey’s post-test. Data are representative of two independent experiments.

### Diltiazem retains its *in vivo* efficacy when administered 24 h after viral infection

To best mimic the therapeutic setting, we next evaluated the efficacy of the same 5-day oral regimen with diltiazem or etilefrine but when initiated 24 h after viral infection (**Figure 4**). As with oseltamivir and monensin, diltiazem treatment completely prevented mortality and reduced weight loss in influenza A(H1N1)pdm09 infected mice, which otherwise showed only 50% (5/10) survival for the etilefrine and saline groups (**Figure 4A-B**). Interestingly, 1-to 1.5-log reductions in LVTs compared to the saline group were observed at day 5 in groups of mice treated with diltiazem or etilefrine (**Figure 4C**). We then used a more stringent approach by increasing the viral inoculum to evaluate the same delayed (24 h post infection) 5-day diltiazem regimen in the context of a 100% lethal A(H1N1)pdm09 infection (**Figure 4D-F**). Whereas treatment with oseltamivir and diltiazem successfully rescued 40% (4/10) and 20% (2/10) of mice, respectively, half-dose treatment with diltiazem (45 mg/kg) rescued 30% (3/10) of mice from death, also showing significant improvement in mean weight loss (**Figure 4D-E**). Calculated hazard ratios (HR) for the saline group compared to these three treatment groups were 8.41 (CI95: 1.65-43.02), 2.85 (0.56-14.47) and 7.62 (1.49-38.96), respectively. Noteworthy, LVTs at day 5 p.i. were comparable among all treated and untreated groups (**Figure 4F**), suggesting mainly a protective effect of diltiazem towards severe influenza infection rather than a direct role in decreasing viral production.

**Fig. 4.**
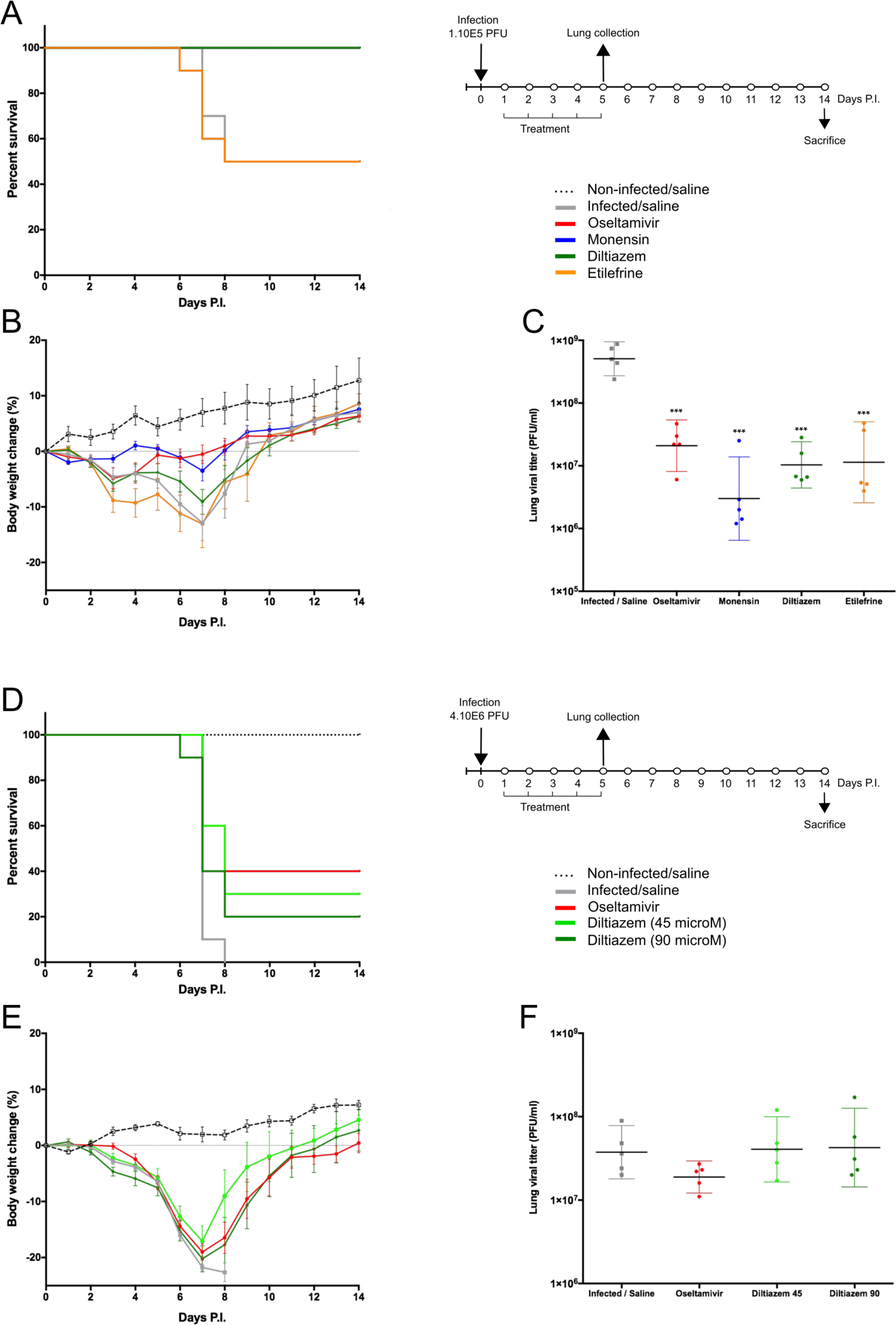
Efficacy of post-infection oral treatment with diltiazem and etilefrine in mice infected with influenza A(H1N1)pdm09 virus. C57BL/6N mice (n=15/group) were intranasally inoculated with 1 x 10^5^ **(A**-**C)** or 4 x 10^6^ **(D**-**F)** PFU of influenza A/Quebec/144147/09 virus on day 0 and treated by gavage with saline (grey), oseltamivir 10 mg/kg/day (red), monensin 10 mg/kg/day (blue, A only), diltiazem 45 mg/kg/day (light green, B only), diltiazem 90 mg/kg/day (dark green), or etilefrine 3 mg/kg/day (orange, A only). A mock-infected, saline-treated group (black dotted line, n=6) was included as control. Treatments were initiated on day 1 (24 h after infection and administered once daily for 5 consecutive days. **(A**, **D)** Survival rates (n=10/group), **(B**, **E)** mean weight changes (±SEM, n=10/group or remaining mice), and **(C**, **F)** median (±CI95, n=5/group) lung viral titers on day 5 p.i. are shown. *p<0.05, **p<0.01 and ***p<0.001 compared to the infected saline-treated group by one-way ANOVA with Tukey’s post-test. Data are representative of two independent experiments.

### Diltiazem significantly reduces viral replication in infected reconstituted human airway epithelia (HAE)

To further complement *in vivo* data, we characterized the inhibitory properties of diltiazem using a biologically relevant reconstituted airway epithelium model, derived from human primary bronchial cells (MucilAir^®^, Epithelix). HAE were infected with influenza A(H1N1)pdm09 at a MOI of 0.1, and treatments on the basolateral medium were initiated 5 h p.i. and continued once daily for 5 consecutive days. In the absence of treatment, viral replication at the apical surface peaked at 48 h p.i. (∼1 x 10^8^ PFU/ml) and was detectable at important levels for at least 7 days. As expected, trans-epithelial electrical resistance (TEER) values, measuring tight junction and cell layer integrity, sharply decreased and bottomed out at 72 h p.i. in the untreated control, correlating with the first virus detection on the basolateral medium (**Figure 5A** and **Table S6**). A similar pattern was observed in infected HAE treated with oseltamivir 0.1 μM or diltiazem 9 μM (CMAP), which conferred no significant advantage over the untreated control. Conversely, oseltamivir 1 μM and diltiazem 90 μM treatments (10-fold CMAP) strongly inhibited viral replication, delaying the peak of viral production by 24 h. Both treatments induced >3-log reductions in apical viral titers at 48 h p.i. compared to the untreated control, and >2-log reductions when comparing peak titers (48 h p.i. untreated vs. 72 h p.i. treated). Moreover, whereas oseltamivir treatment stabilized TEER during the time-course of infection, diltiazem treatment partially buffered the TEER decrease observed in the untreated control (**Figure 5A** and **Table S6**). No virus was detected on the basolateral medium for these two treated groups, and absence of treatment-induced toxicity was confirmed by measuring the release of intracellular lactate dehydrogenase (LDH). Interestingly, we observed that inhibitory and protective properties demonstrated by diltiazem were progressively reversible when basolateral medium was replaced with fresh medium without drugs. Overall, these results are in accordance and strongly support the inhibitory and protective effects of diltiazem observed *in vitro* and in mice, respectively.

**Fig. 5.**
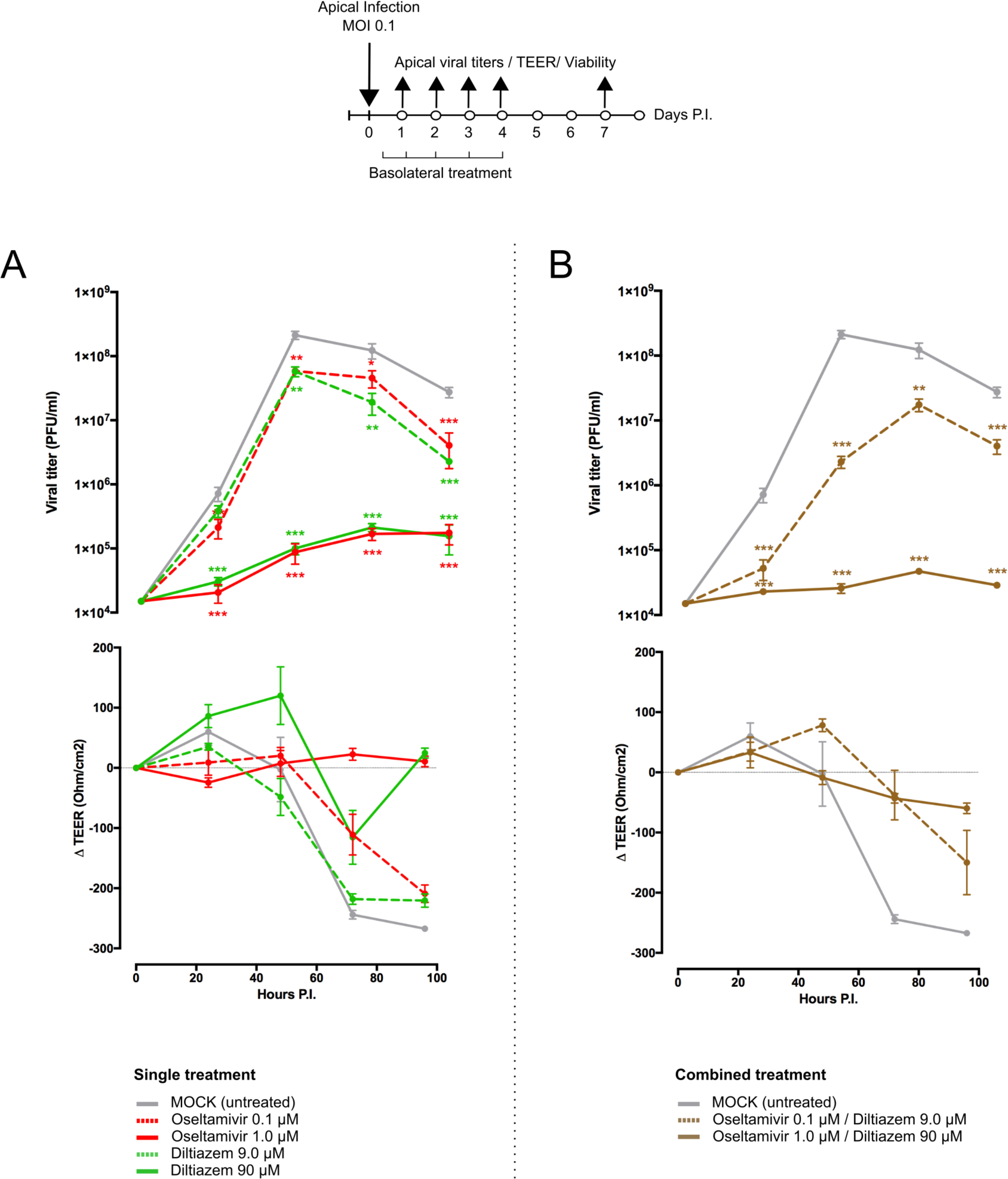
Diltiazem significantly reduces viral replication in infected reconstituted human airway epithelia (HAE). Apical viral production (±SEM) and transepithelial electrical resistance (Δ TEER±SEM) in MucilAir^®^ human airway epithelium infected on the apical pole with influenza A/Lyon/969/09 (H1N1)pdm09 virus at a MOI of 0.1 and subjected to **(A)** single or **(B)** combined treatments by the basolateral pole. Treatments with culture medium (mock, grey), oseltamivir 0.1 µM (red, dotted line), oseltamivir 1 µM (red, solid line), diltiazem 9 µM (green, dotted line), diltiazem 90 µM (green, solid line), oseltamivir 0.1 µM / diltiazem 9 µM (brown, dotted line) or oseltamivir 1 µM / diltiazem 90 µM (brown, solid line) were initiated 5 h after infection and administered once daily for 5 consecutive days. *p<0.05, **p<0.01 and ***p<0.001 compared to the infected mock-treated group by one-way ANOVA with Tukey’s post-test. Data are representative of three independent experiments.

### Diltiazem-oseltamivir combination confers improved efficacy when compared to monotherapy in infected HAE

We anticipated that the combination of two antiviral compounds that target different viral/cellular determinants could induce better virological and physiological responses when compared to antiviral monotherapy. We therefore evaluated the diltiazem-oseltamivir combination in the same conditions described above, notably a 5-day treatment course with treatment initiation at 5 h p.i. The diltiazem 90 μM / oseltamivir 1 μM combination conferred >3-log reduction in apical peak viral titers when compared to the untreated control, even greater than that observed with same dose monotherapy. TEER values remained stable during combined treatment, comparable to those observed with oseltamivir 1 μM monotherapy (**Figure 5B** and **Table S6**). Remarkably, although not effective as monotherapy in the low concentrations tested above, the diltiazem 9 μM / oseltamivir 0.1 μM combination contrariwise delayed the peak of viral production, significantly reduced apical viral titers, and slightly buffered TEER values compared to the untreated control (**Figure 5B** and **Table S6**). Once again, no treatment-related toxicity was observed for any of the combinations tested. These results plead in favor of the potential of diltiazem for the improvement of current anti-influenza therapy with neuraminidase inhibitors.

### Diltiazem treatment induces a significant reversion of the viral infection signature

Since the rationale behind our approach relies on attaining antiviral activity through a drug-induced global and multi-level inversion of the infection signature, we advantageously used the MucilAir^®^ HAE model coupled with high-throughput sequencing in order to characterize and compare the specific transcriptional signatures induced by infection and/or diltiazem treatment (**Figures 6 and S4**). HAE were mock-infected or infected with influenza A(H1N1)pdm09 virus and then mock-treated or treated in the same experimental conditions in which the antiviral effect of diltiazem has been previously validated (MOI of 0.1, 90 μM diltiazem). At 72 h p.i., cells were lysed and total RNA was extracted. cDNA libraries were then produced, amplified, and subjected to high-throughput sequencing. Taking the mock-infected / mock-treated (“mock”) as baseline, we initially performed DAVID functional gene enrichment (absolute fold change >2, Benjamini-Hochberg corrected p-value <0.05) on the specific transcriptional signature of diltiazem with the objective of gaining insight on the putative host pathways involved in its antiviral effect. The lists of up-regulated (n=194) and down-regulated (n=110) transcripts in the mock-infected / diltiazem (“mock + diltiazem”) condition were analyzed using DAVID 6.8 to highlight associations with specific GO terms. Although no enriched BP was identified among down-regulated transcripts, the list of up-regulated transcripts associated with diltiazem treatment highlighted 7 particularly enriched BP. While 4 of these BP (GO:0009615; GO:0045071; GO:0051607; GO:0060337) are directly linked to antiviral response/cellular response to virus, the remaining 3 (GO:0055114; GO:0008299; GO:0006695) are involved in cholesterol biosynthesis/metabolism (**Figure 6A**). We then compared the common differentially expressed transcript levels between the three infection/treatment conditions. These transcriptional signatures revealed a marked anti-correlated profile between the “mock + diltiazem” and the infected / mock-treated (“H1N1”) conditions (**Figure 6B**), supported by a median Spearman’s ρ correlation value of -0.82 (**Figure 6C**). Most important, the infected / diltiazem (“H1N1 + diltiazem”) condition yielded ρ correlation values of 0.40 and -0.72 when compared to either “mock + diltiazem” or “H1N1”, respectively, therefore confirming a partial reversion of the infection virogenomic signature during effective antiviral treatment with diltiazem (**Figures 6D and S4**), as expected.

**Fig. 6.**
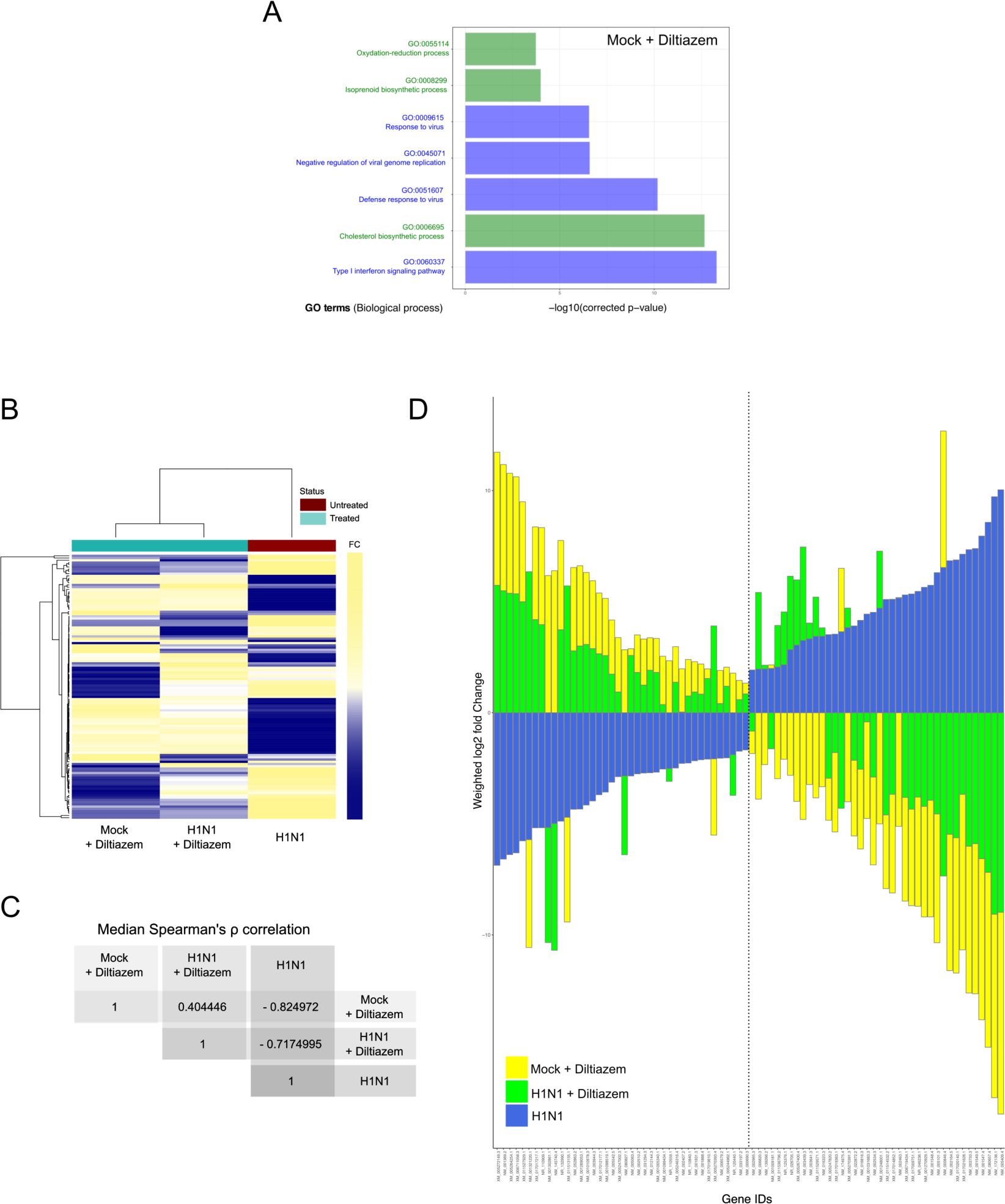
Diltiazem treatment effectively induces significant reversion of the viral infection signature. **(A)** DAVID gene enrichment analysis of the diltiazem transcriptional signature. The seven most significant biological processes (BP) are presented. BP related to antiviral response and cholesterol biosynthesis/metabolism are represented in blue and green, respectively. **(B)** Hierarchical clustering and heatmap of the 118 common differentially expressed transcripts (absolute fold change >2, Benjamini-Hochberg corrected p-value <0.05) between mock-infected / diltiazem (“mock + diltiazem”), infected / mock-treated (“H1N1”), or infected / diltiazem (“H1N1 + diltiazem”) HAE. The mock-infected / mock-treated (“mock”) condition was used as baseline. Mean-weighted fold changes are color-coded from blue to yellow. **(C)** Median Spearman ρ correlation value calculations between the 3 conditions highlighted in the heatmap. **(D)** Stacked barplot representation of the 40 most up/down-regulated transcripts highlighted in the analysis. Barplots were constructed in R3.3.1 based on mean-weighted fold changes and ordered according to H1N1 values (blue). Mock + diltiazem and H1N1 + diltiazem conditions are represented in yellow and green, respectively.

## Discussion

The existing urge for alternative strategies to cope with the limited efficacy of currently approved antivirals for the prevention and treatment of influenza infections [2, 41, 42], mostly in the case of patients with severe influenza and acute respiratory distress syndrome (ARDS) [43, 44], represented the central driving force of this study. Here, we developed and validated for the first time an innovative approach based on clinical genomic signatures of respiratory viral infections for the rapid discovery, *in vitro, in vivo* and *ex-vivo* evaluation, as well as the repurposing of FDA-approved drugs for their newly identified host-targeted inhibitory and protective properties against influenza infections.

Targeting host components on which viral replication depends instead of viral determinants represents a real change of paradigm in antiviral development, with pioneering results mainly observed in the context of antiretroviral therapy [13, 45]. Nevertheless, and despite strong putative advantages such as the achievement of broad-spectrum antiviral efficacy and the minimization of viral drug resistance, this approach usually fails to overcome two major limiting factors of classic compound screening. Firstly, it remains target-centered *per se*, therefore leading to the identification of drugs with limited efficacy due to the complex network and high redundancy of the host cellular pathways. Secondly, the need of high-throughput screenings often entails the measurement of a very limited number of viral parameters, usually in non-physiologically and hence poorly relevant conditions and/or cellular models.

Based on our initial proof-of-concept study on the *in silico* screening of the CMAP database [19, 20] with no initial *a priori* on specific host targets [18], we moved our approach up to the clinical trial setting, by determining exploitable and more relevant virogenomic profiles directly from standard clinical samples of influenza-infected patients. Since the low amount of often degraded RNA obtained from these samples represented a major challenge, we implemented an original combination of sample preparation techniques for low input but high quality samples with data processing initially designed for expression analysis of non-model species [22, 23].

Another substantial development was the integration of several lists of candidate molecules issued from different transcriptomic signatures with enriched relevant DAVID Gene Ontology terms, and their final selection based on their pharmacological classes and potential compatibility as antivirals. Our refined strategy allowed the selection of a shortlist of 35 high potential candidates out of a rationalized computational screening of a total of 1,309 FDA-approved bioactive molecules. This drastic positive selection step constituted a major advantage, since it enabled the implementation of relevant and integrated *in vitro, in vivo* and *ex-vivo* evaluations in a time- and cost-effective manner. Most important, the use of patient (*in vivo)* virogenomic profiles led to the identification of molecules with highly improved *in vitro* activity and significant *in vivo* antiviral efficacy as compared with compounds previously obtained from our initial study based on cell culture (*in vitro)* virogenomic profiles [18]. These results truly highlight the added value of using relevant clinical virogenomic signatures to optimize the computational screening for active drugs.

Two of the molecules identified in this study with transcriptomic profiles that counteract clinical virogenomic signatures (e.g. ribavirin and monensin) have already been validated for their anti-influenza properties [38, 40], and then supported the relevance of our compound selection strategy. Nevertheless, although different modes of action have been postulated for the anti-influenza activity of the synthetic guanosine analog ribavirin [39], the exact mechanisms remain uncharacterized so far. Similarly, it has been postulated that monensin, an antibiotic isolated from *Streptomyces spp*, may have a role as a ionophore that interferes with intracellular transport of several enveloped viruses, including influenza [40]. In that sense, even if we cannot rule out that some of the molecules identified *in silico* exert a direct effect on a specific pathway or cellular target, the fact that these molecules have been identified with a high anti-correlation rate in CMAP strongly supports a potential multi-target inhibitory effect, probably resulting in deep modifications of host gene expression. In fact, both monensin and ribavirin were previously reported to modulate the host cellular gene expression profile, notably through the up-regulation of the cholesterol and lipid biosynthesis genes [46] or the virus-induced ISRE signaling and antiviral ISGs genes [47], respectively.

The two most promising molecules highlighted in this study are etilefrine, an alpha and beta-adrenergic receptor agonist, currently indicated as a cardiotonic and anti-hypotensive agent [48] and mainly diltiazem, a voltage-gated Ca2+ channel antagonist that is currently used to control angina pectoris and cardiac arrhythmia [49]. In addition to their strong inhibitory effect on the viral growth of circulating A(H1N1)pdm09 viruses, with *in vitro* EC50 values in the nanomolar range (**Figure 2**), both molecules also demonstrated antiviral properties against oseltamivir-resistant A(H1N1)pdm09 and prototype H3N2 and B influenza strains (**Table S4**). Our *in vivo* results (**Figures 3**-**4**), obtained without previous treatment optimization in terms of dosage or administration route, also suggest that these drugs harbor a protective role towards influenza infection, particularly in the case of diltiazem, which conferred increased survival in mice even in a model of severe influenza infection (**Figure 4D**-**F**). Moreover, the inhibitory and protective properties of diltiazem were validated in the reconstituted human airway epithelium model, also showing enhanced efficacy when combined with oseltamivir (**Figure 5**).

Finally, a very recent study by Fujioka and colleagues [50] confirmed the antiviral activity of diltiazem anticipated by our approach. In that study, based on the role of Ca2+ channels on the attachment of influenza viruses to the host cell, the authors discuss whether the diltiazem induced modulation of Ca2+ channel activity might not fully explain such observed antiviral activity, consistent with a multi-level (off-target) effect of diltiazem. In this context, in which not all Ca2+ channel inhibitors confer significant antiviral activity, the newly described capacity of diltiazem to partially reverse the global virogenomic signature of infection and modulate specific genes related to the host antiviral response and cholesterol metabolism (**Figures 6 and S4**) suggests a putative explanation for its inhibitory effect observed *in vitro, ex vivo* and in mice. Nevertheless, further investigations are underscored to shed light on the specific mechanisms underlying such potential multi-level mode of action of diltiazem.

## Conclusions

Overall, the results presented here set a solid baseline for our drug repurposing strategy and for the use of diltiazem as a host-targeted antiviral in clinical practice. Moreover, the increased antiviral efficacy observed in reconstituted human airway epithelium (**Figures 5B** and **S6 Table**) plead in favor of the combination of diltiazem with the virus-targeted antiviral oseltamivir for the improvement of current anti-influenza therapy, and possibly decreasing the risk of development of viral resistance. In that regard, our results prompted a French multicenter randomized clinical trial aimed at assessing the effect of diltiazem-oseltamivir bitherapy compared with standard oseltamivir monotherapy for the treatment of severe influenza infections in intensive care units, hence completing the bedside-to-bench and bench-to-bedside cycle of our innovative approach. Additionally, retrospective signature analysis of sequential respiratory samples from patients included in both study arms and stratified according to their clinical response to treatment will provide valuable data to pursue the investigations on the specific mediators of the diltiazem-related antiviral response. This trial (FLUNEXT TRIAL PHRC #15-0442, ClinicalTrials.gov identifier NCT03212716) is currently ongoing.

Finally, our study underscores the high value of clinical specimens and the advantages of exploiting virogenomic and chemogenomic data for the successful systematic repurposing of drugs already available in our modern pharmacopeia as new effective antivirals. We propose that our approach targeting respiratory epithelial cells, the principal influenza infected cell type in the lung, could be extended to other respiratory viruses and eventually to other pathogens involved in acute infections. Importantly, drug repurposing presents several financial and regulatory advantages compared to the development of *de novo* molecules [5], which are of particular interest not only in the context of antimicrobial resistance but also against both emerging or recurrent pathogens for which we are still disarmed.

## Declarations

### Ethics approval and consent to participate

Adult patients were recruited by general practitioners in the context of a previously published randomized clinical trial (Escuret et al., 2012) (ClinicalTrials.gov identifier NCT00830323) and all of them provided written informed consent. The study protocol was approved by the Lyon Ethics Committee (Comité de Protection des Personnes Lyon B) on September 9th, 2009 and conducted in accordance with the Declaration of Helsinki.

All animal procedures were approved by the Institutional Animal Care Committee of the Centre Hospitalier Universitaire de Québec (CPAC protocol authorization #2012-068-3) according to the guidelines of the Canadian Council on Animal Care.

### Consent for publication

Not applicable.

### Availability of data and material

The main datasets supporting the conclusions of this article are included within the article and its additional files. All raw microarray data from virogenomic signatures were deposited on the National Center for Biotechnology Information’s Gene Expression Omnibus (GEO), under accession number GSE93731. Other datasets used and/or analyzed during the current study are available from the corresponding authors on reasonable request.

### Authors’ contributions

AP, OT, JT, GB and MRC designed and coordinated the study. VE, JP and BL acquired clinical data from the cohort. AP, OT, TJ, BP, AT, MR, MEH, CR, CNL, SC, CLL and JT performed experiments and provided technical support. AP, OT, MR, JP, JT, GB and MRC interpreted data. AP, OT and MRC wrote the manuscript. JT, GB and MRC revised the manuscript.

## Acknowledgements

The authors want to thank Jacques Corbeil and Frederic Raymond (Research Center in Infectious Diseases of the CHU de Quebec and Laval University, Quebec) for their help and useful advice.

## Supplementary Materials

**Fig. S1.**
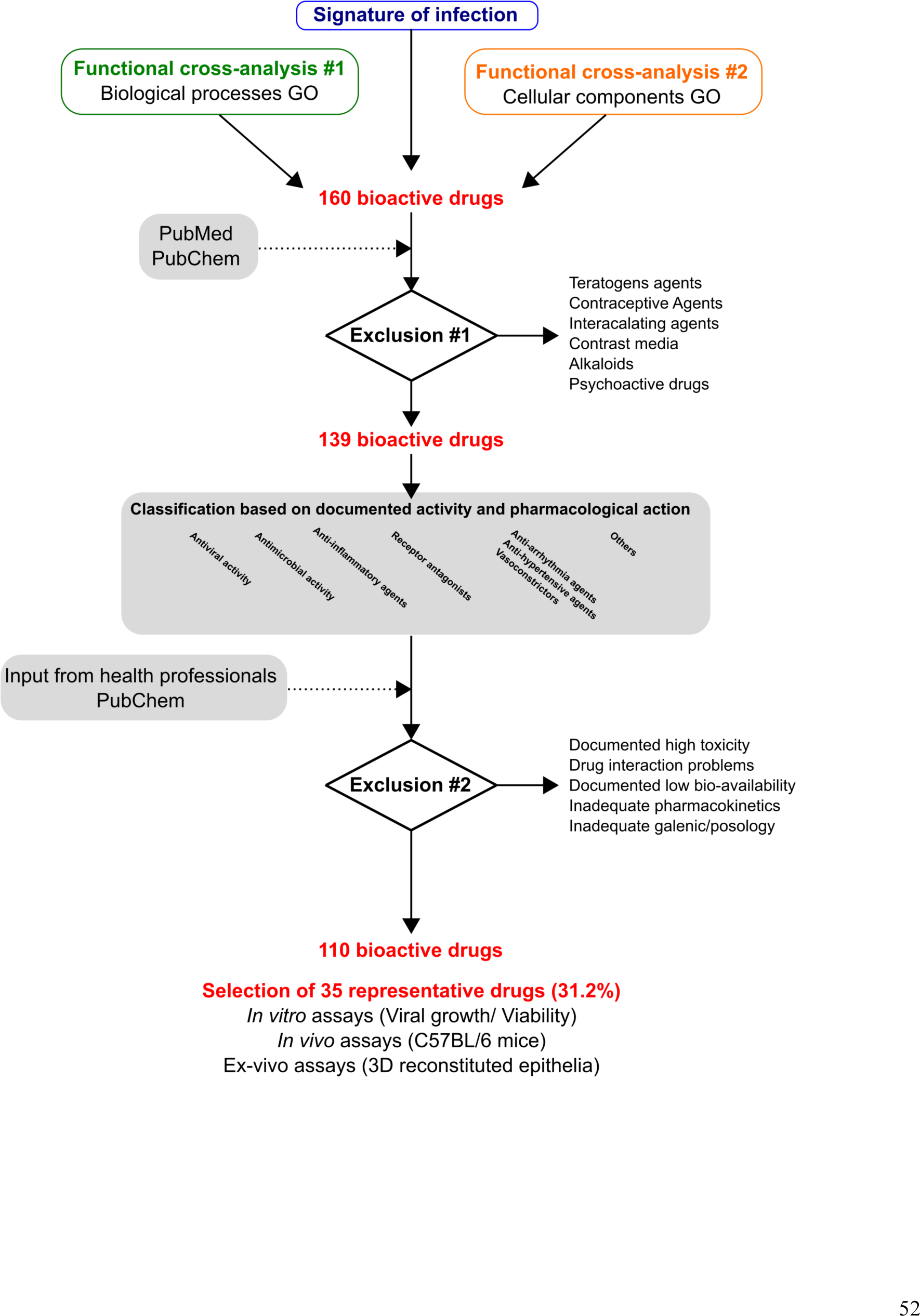
Decision tree used to rationally reduce the number of drug candidates. Bioactive molecules were excluded if not compatible with a final use as antiviral, mostly for safety (e.g. teratogens, intercalating agents) and/or pharmacological (e.g. documented low bioavailability) reasons. An additional selection level based on analysis of documented pharmacological actions was included, to finally define a shortlist of 35 representative molecules (<3% of CMAP) for in vitro screening (Table 1).

**Fig. S2.**
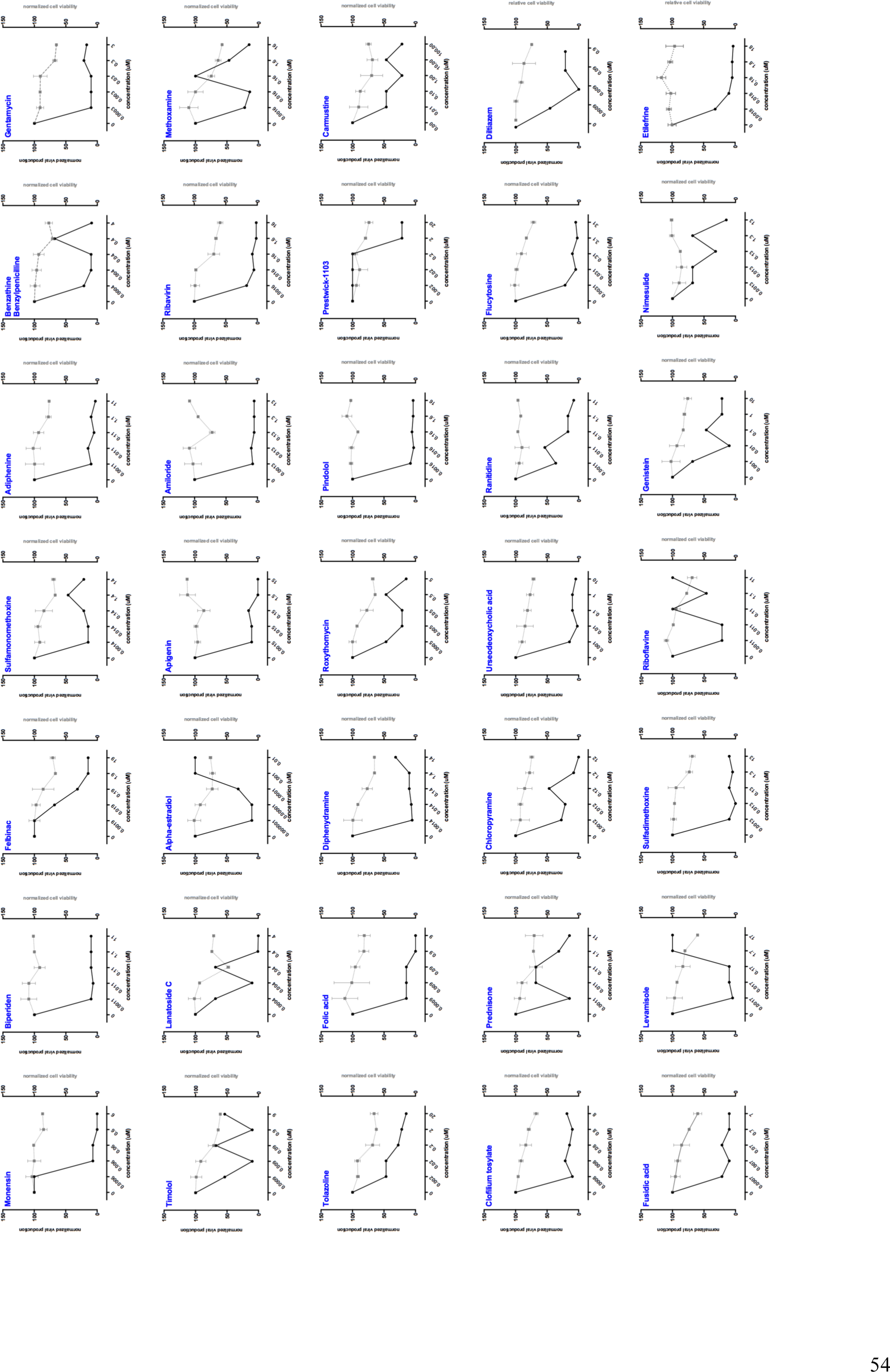
Dose-response curves for the 35 molecules tested *in vitro*. In comparison with a mock-treated control, the impact of pre-treatment/treatment on % relative viral production (black line, left Y axis) and % relative cell viability (grey line, right Y axis) was measured at the indicated concentrations.

**Fig. S3.**
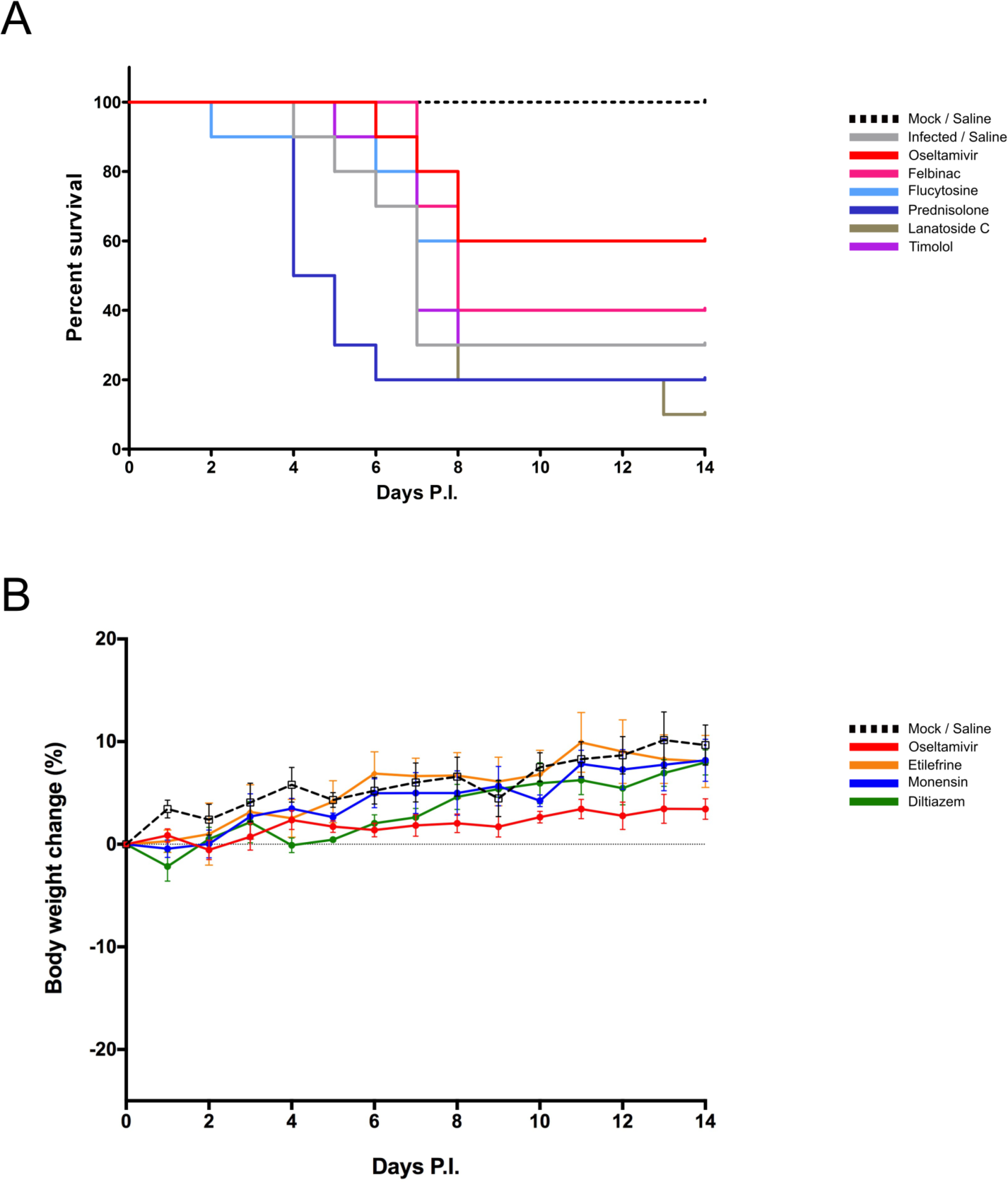
Efficacy and toxicity after oral administration of selected molecules in mice. (A) Survival curves of C57BL/6N mice (n=15/group), intranasally inoculated with 5 x 10^5^ PFU of influenza A/Quebec/144147/09 virus on day 0 and treated by gavage with saline (grey), oseltamivir 10 mg/kg/day (red), lanatoside C 100 mg/kg/day (olive), prednisolone 5 mg/kg/day (dark blue), flucytosine 240 mg/kg/day (light blue), felbinac 5 mg/kg/day (fuchsia) or timolol 50 mg/kg/day (purple). A mock-infected, saline-treated group (black dotted line, n=6) was included as control. Treatments were initiated on day 0 (6 h before infection) and administered once daily for 5 consecutive days. (B) Body weight changes of mock-infected C57BL/6N mice (n=10/group) treated by gavage with saline (grey), oseltamivir 10 mg/kg/day (red), monensin 10 mg/kg/day (blue), diltiazem 90 mg/kg/day (green), or etilefrine 3 mg/kg/day (orange). A saline-treated group (black dotted line, n=6) was included as control. Treatments were initiated on day 0 (6 h before mock-infection) and administered once daily for 5 consecutive days.

**Fig. S4.**
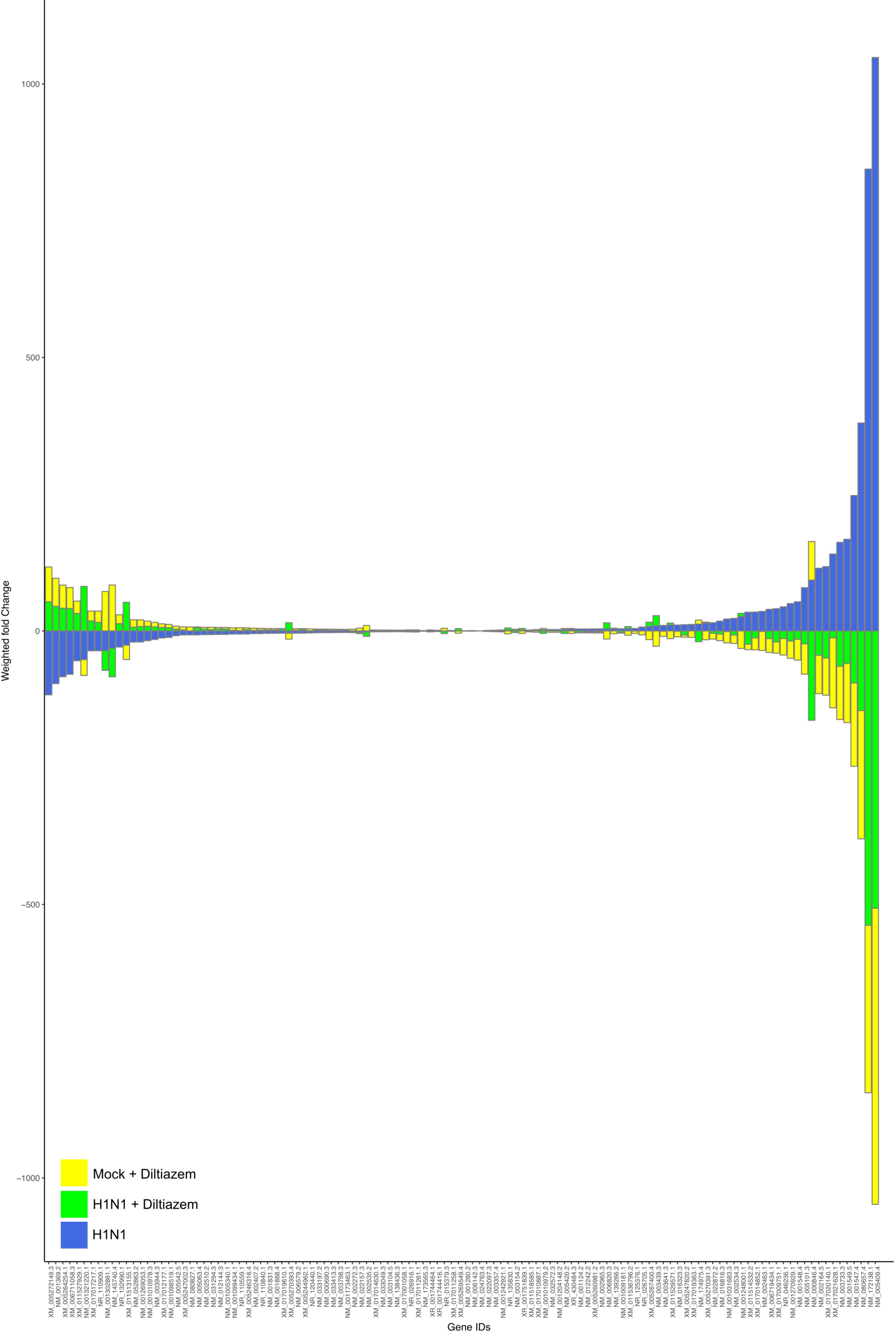
Diltiazem treatment effectively induces significant reversion of the viral infection signature. Stacked barplots with mean-weighted fold changes for the complete list of the 118 common differentially expressed transcripts (absolute fold change >2, Benjamini-Hochberg corrected p-value <0.05) between the mock + diltiazem (yellow), H1N1 (blue), and H1N1 + diltiazem (green) conditions. Barplots were constructed in R3.3.1 based on mean-weighted fold changes and ordered according to H1N1 values (blue).

**Table S1.**
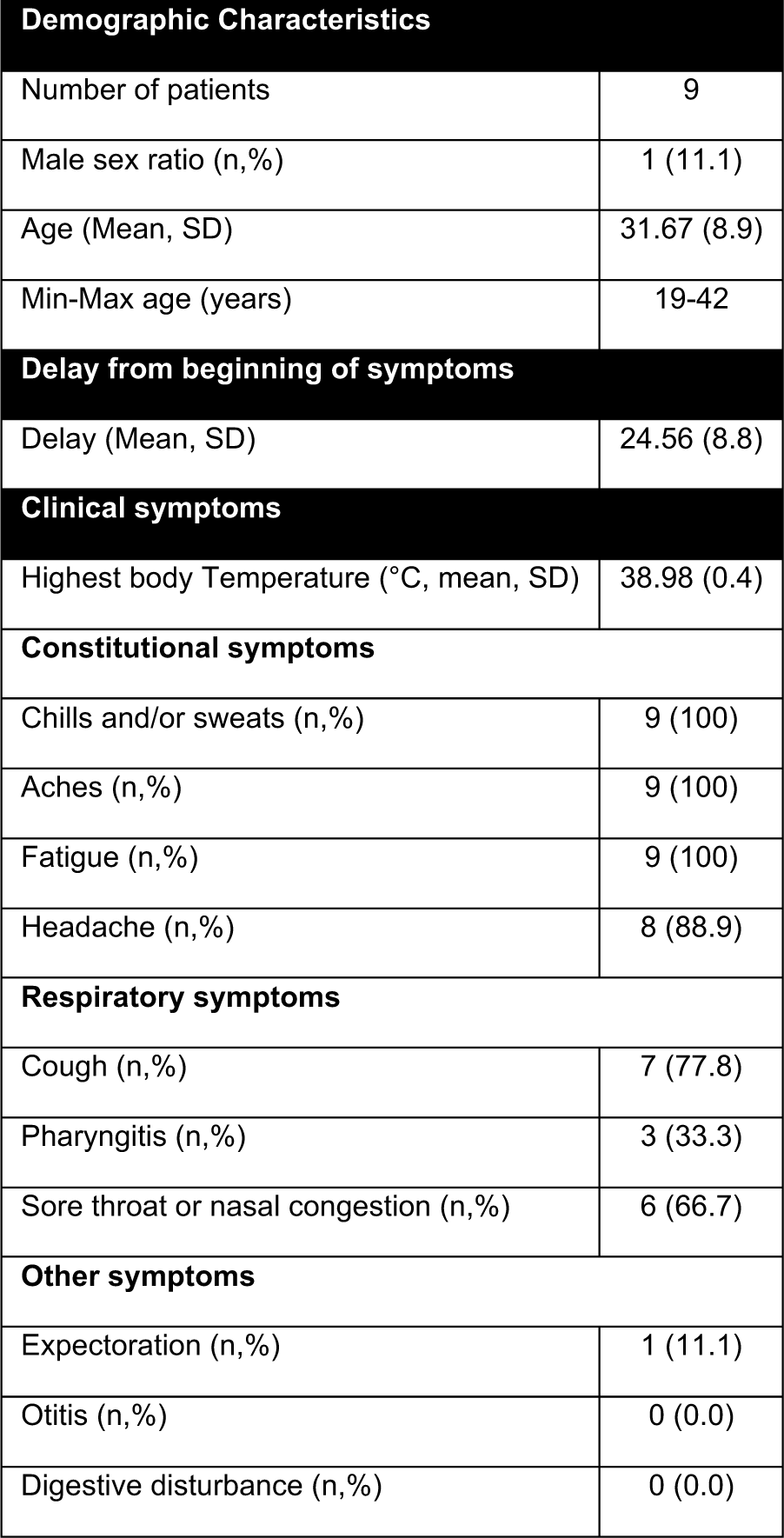
Demographic and clinical characteristics of patients included in the study. Samples were obtained in the context of a previous clinical trial conducted in France during the A(H1N1)pdm09 pandemic, aimed at evaluating the antiviral efficacy and tolerability of classic antiviral monotherapy versus bitherapy (Escuret et al., 2012).

**Table S2.** **List of 160 selected molecules and their documented pharmacological classes.** The 35 selected compounds for *in vitro* and *in vivo* evaluation are highlighted in grey. Asterisks (*) indicate molecules previously evaluated for their antiviral properties against influenza viruses or other viruses according to the literature, and question marks (?) indicate absence of assigned pharmacological class. Documented pharmacological classes were obtained from PubChem (https://pubchem.ncbi.nlm.nih.gov).

| Molecules | Pharmalogical class |
| --- | --- |
| Monensin* | Antifungal Agents/Antiprotozoal Agents/Coccidiostats/Proton Ionophores/Sodium Ionophores |
| Isoxicam | Anti-Inflammatory Agents, Non-Steroidal |
| Iloprost* | Platelet Aggregation Inhibitors/Vasodilator Agents |
| Securinine | Alkaloids/GABAA Receptor Antagonists |
| Prestwick-692 | Steroid Alkaloids |
| Biperiden* | Antiparkinson Agents/Muscarinic Antagonists/Parasympatholytics |
| Clorsulon | Anthelmintics/Antiplatyhelminic Agents |
| Felbinac | Anti-Inflammatory Agents, Non-Steroidal |
| Chenodeoxycholic acid* | Cathartics/Gastrointestinal Agents |
| Sulfamonomethoxine | Anti-Infective Agents |
| 3-acetamidocoumarin | ? |
| Meteneprost | Abortifacient Agents, Nonsteroidal |
| Adiphenine | Parasympatholytics/Anticholinergics/Antispasmodics |
| Benzathine benzylpenicillin | Anti-Bacterial Agents |
| Gentamicin | Anti-Bacterial Agents/Protein Synthesis Inhibitors |
| Timolol* | Adrenergic beta-Antagonists/Anti-Arrhythmia Agents/Antihypertensive Agents |
| Atractyloside* | Enzyme Inhibitors |
| Nadolol | Adrenergic beta-Antagonists/Anti-Arrhythmia Agents/Antihypertensive Agents/Sympatholytics |
| Lycorine* | Alkaloids/Protein Synthesis Inhibitors |
| Tranexamic acid* | Antifibrinolytic Agents |
| Lanatoside C | Anti-Arrhythmia Agents |
| Podophyllotoxin* | Antineoplastic Agents, Phytogenic/Keratolytic Agents/Tubulin Modulators |
| Alpha-estradiol* | 5alpha-reductase inhibitors/Androgenic alopecia treatment |
| Alprostadiol | Fibrinolytic Agents/Platelet Aggregation Inhibitors/Vasodilator Agents |
| Apigenin* | Anti-Inflammatory Agents, Non-Steroidal/? |
| Pipenzolate bromide | Muscarinic Receptor Antagonists |
| Amiloride* | Epithelial Sodium Channel Blockers/Diuretics/Acid Sensing Ion Channel Blockers |
| STOCK1N-35696 | ? |
| Carmustine | Antineoplastic Agents, Alkylating |
| PF-00539745-00 | ? |
| Clofilium tosylate | Anti-Arrhythmia Agents |
| Prestwick-983* | Antimetabolites/Antiviral Agents |
| Prestwick-675 | Anthelmintics/Anticestodal Agents/Antiprotozoal Agents/Tubulin Modulators |
| Atracurium besilate | Neuromuscular Nondepolarizing Agents/Nicotinic Antagonists |
| Xamoterol | Adrenergic beta-1 Receptor Agonists |
| Demecarium bromide | Cholinesterase Inhibitors |
| Iopromide | Contrast Media |
| Etilefrine | Adrenergic alpha-Agonists/Adrenergic beta-1 Receptor Agonists/Cardiotonic Agents/Sympathomimetics/Vasoconstrictor Agents |
| Iproniazid | Antidepressive Agents/Monoamine Oxidase Inhibitors |
| Ketotifen* | Anti-Allergic Agents/Antipruritics/Histamine H1 Antagonists |
| Gly-His-Lys | Anti-Inflammatory Agents, Non-Steroidal? |
| Canadine* | Anti-Arrhythmia Agents/Calcium Channel Blockers/Platelet Aggregation Inhibitors |
| Viomycin | Anti-Bacterial Agents/Antibiotics, Antitubercular/Protein Synthesis Inhibitors |
| **Disopyramide** | Anti-Arrhythmia Agents/Voltage-Gated Sodium Channel Blockers |
| **Fusidic acid** | Anti-Bacterial Agents/Protein Synthesis Inhibitors |
| **Amiprilose** | Adjuvants, Immunologic/Anti-Inflammatory Agents, Non-Steroidal/Antiviral Agents |
| **Anisomycin\*** | Anti-Bacterial Agents/Antiprotozoal Agents/Nucleic Acid Synthesis Inhibitors/Protein Synthesis Inhibitors |
| **Ajmaline** | Anti-Arrhythmia Agents |
| **Arecoline** | Cholinergic Agonists |
| **Metampicillin** | Anti-Bacterial Agents |
| **Lasalocid** | Anti-Bacterial Agents/Coccidiostats/Ionophores |
| **Tolnaftate** | Antifungal Agents |
| **Metixene** | Muscarinic Receptor Antagonists |
| **PF-00539758-00** | ? |
| **Streptomycin** | Anti-Bacterial Agents/Protein Synthesis Inhibitors |
| **Sulfapyridine** | Anti-Infective Agents/Dermatologic Agents |
| **Pivmecillinam** | Anti-Bacterial Agents/Anti-Infective Agents, Urinary |
| **Ribavirin\*** | Antimetabolites/Antiviral Agents |
| **Prestwick-642\*** | Dermatologic Agents/Teratogens |
| **Bumetanide** | Diuretics/Sodium Potassium Chloride Symporter Inhibitors |
| **Prednisone** | Anti-Inflammatory Agents/Antineoplastic Agents, Hormonal/Glucocorticoids |
| **Doxylamine** | Antiemetics/Histamine H1 Antagonists |
| **Diphenylpyraline** | Histamine H1 Antagonists |
| **Finasteride** | 5-alpha Reductase Inhibitors |
| **Rosiglitazone\*** | Hypoglycemic Agents |
| **15-delta prostaglandin J2** | Anti-Inflammatory Agents, Non-Steroidal?/NFkB inhibitor? |
| **5230742** | ? |
| **Chloropyramine** | Histamine H1 Antagonists |
| **Prestwick-685** | Anti-Inflammatory Agents, Non-Steroidal/Coloring Agents/Leprostatic Agents |
| **Nimesulide\*** | Anti-Inflammatory Agents, Non-Steroidal/Cyclooxygenase Inhibitors |
| **Morantel** | Anthelmintics/Antinematodal Agents |
| **Tropicamide** | Muscarinic Receptor Antagonists/Mydriatics |
| **Piperidolate** | Muscarinic Receptor Antagonists |
| **Riboflavin\*** | Photosensitizing Agents/Vitamin B Complex |
| **Methoxamine** | Adrenergic alpha-1 Receptor Agonists/Sympathomimetics/Vasoconstrictor Agents |
| **Hydrocotarnine** | Alkaloids/? |
| **Propidium iodide** | Coloring Agents/Indicators and Reagents/Intercalating Agents |
| **Tolazoline** | Adrenergic alpha-Antagonists/Antihypertensive Agents/Vasodilator Agents |
| **5279552** | ? |
| **Sulfadimethoxine** | Anti-Infective Agents |
| **N-acetyl-L-leucine** | *Vertigo treatment/?* |
| **Gibberellic acid** | Plant Growth Regulators |
| **Clorgiline** | Antidepressive Agents/Monoamine Oxidase Inhibitors |
| **Genistein** | Anticarcinogenic Agents/Phytoestrogens/Protein Kinase Inhibitors |
| **Pargyline** | Antihypertensive Agents/Monoamine Oxidase Inhibitors |
| **Cortisone** | Anti-Inflammatory Agents |
| **Medrysone** | Anti-inflammatory Agents/Glucocorticoids |
| **Isoflupredone** | Anti-inflammatory Agents/Mineralcorticoids |
| **Prestwick-1082** | ? |
| **Aciclovir\*** | Antiviral Agents |
| **Sulconazole** | Antifungal Agents |
| **Cycloserine** | Anti-Infective Agents, Urinary/Antibiotics, Antitubercular/Antimetabolites |
| **Procainamide** | Anti-Arrhythmia Agents/Voltage-Gated Sodium Channel Blockers |
| **Chlortalidone** | Antihypertensive Agents/Diuretics/Sodium Chloride Symporter Inhibitors |
| **Chlorzoxazone** | Muscle Relaxants, Central |
| **Oxolamine** | Antitussive Agents |
| **Folic acid** | Hematinics/Vitamin B Complex |
| **Furazolidone** | Anti-Infective Agents, Local/Anti-Infective Agents, Urinary/Antitrichomonal Agents/Monoamine Oxidase Inhibitors |
| **Cotinine** | Indicators and Reagents |
| **Ikarugamycin\*** | Anti-Infective Agents |
| **H-7** | Enzyme Inhibitors |
| **Natamycin** | Anti-Bacterial Agents/Anti-Infective Agents, Local/Antifungal Agents |
| **H-89\*** | Protein Kinase Inhibitors |
| **Guanadrel** | Antihypertensive Agents |
| **Midodrine** | Adrenergic alpha-1 Receptor Agonists/Sympathomimetics/Vasoconstrictor Agents |
| **Etiocholanolone** | Ketosteroids |
| **Methyldopate** | Antihypertensive Agents |
| **Oxymetazoline\*** | Adrenergic alpha-Agonists/Nasal Decongestants/Sympathomimetics |
| **Levomepromazine** | Analgesics, Non-Narcotic/Antipsychotic Agents/Dopamine Antagonists |
| **Thapsigargin** | Enzyme Inhibitors |
| **Pyrithyldione** | Psychoactive drugs |
| **Nicergoline** | Adrenergic alpha-Antagonists/Nootropic Agents/Vasodilator Agents |
| **Apramycin** | Anti-Bacterial Agents |
| **Prestwick-1103** | Anti-Inflammatory Agents, Non-Steroidal/Cyclooxygenase Inhibitors |
| **Fenoprofen** | Anti-Inflammatory Agents, Non-Steroidal/Cyclooxygenase Inhibitors |
| **Fludrocortisone** | Mineralocorticoids/Anti-Inflammatory Agents |
| **Diphenhydramine** | Anesthetics, Local/Anti-Allergic Agents/Antiemetics/Histamine H1 Antagonists/Hypnotics and Sedatives |
| **Naloxone** | Narcotic Antagonists |
| **Benzonatate** | Antitussive Agents |
| **Thiocolchicoside** | Muscle Relaxants |
| **Eucatropine** | Mydriatics |
| **Dextromethorphan** | Antitussive Agents/Excitatory Amino Acid Antagonists |
| **Isometheptene** | Adrenergic alpha-1 Receptor Agonists/Sympathomimetics/Vasoconstrictor Agents |
| **Cinoxacin** | Anti-Infective Agents |
| **Levamisole\*** | Adjuvants, Immunologic/Antinematodal Agents/Antirheumatic Agents |
| **Ursodeoxycholic acid** | Cholagogues and Cholaretics |
| **4,5-dianilinophthalimide** | Protein Kinase Inhibitors |
| **Ifenprodil** | Adrenergic alpha-Antagonists/Excitatory Amino Acid Antagonists/Vasodilator Agents |
| **CP-320650-01** | ? |
| **Roxithromycin** | Anti-Bacterial Agents |
| **Lisuride** | Antiparkinson Agents/Dopamine Agonists/Serotonin Receptor Agonists |
| **lomefloxacin** | Anti-Infective Agents |
| **Iorglumide** | Hormone Antagonists |
| **Piretanide** | Diuretics/Sodium Potassium Chloride Symporter Inhibitors |
| **L-methionine sulfoximine\*** | Enzyme Inhibitors |
| **Diltiazem** | Antihypertensive Agents/Calcium Channel Blockers/Cardiovascular Agents/Vasodilator Agents |
| **Tyloxapol** | Detergents/Surface-Active Agents |
| **Flumequine** | Anti-Infective Agents/Anti-Infective Agents, Urinary |
| **Terazosin** | Adrenergic alpha-1 Receptor Antagonists |
| **Triflusal** | Platelet Aggregation Inhibitors |
| **Ranitidine** | Anti-Ulcer Agents/Histamine H2 Antagonists |
| **Flucytosine\*** | Antifungal Agents/Antimetabolites |
| **Etomidate** | Anesthetics, Intravenous/Hypnotics and Sedatives |
| **Dioxybenzone** | UVB/UVA protection? |
| **Furaltadone** | Anti-Infective Agents, Urinary |
| **Ornidazole** | Amebicides/Antitrichomonal Agents/Radiation-Sensitizing Agents |
| **Dicloxacillin** | Anti-Bacterial Agents |
| **Pindolol** | Adrenergic beta-Antagonists/Antihypertensive Agents/Serotonin Antagonists/Vasodilator Agents |
| **Tretinoin\*** | Antineoplastic Agents/Keratolytic Agents |
| **Proscillaridin** | Cardiotonic Agents/Enzyme Inhibitors |
| **Ouabain\*** | Cardiotonic Agents/Enzyme Inhibitors |
| **Beclometasone** | Anti-Asthmatic Agents/Anti-Inflammatory Agents/Glucocorticoids |
| **Mexiletine** | Anti-Arrhythmia Agents/Voltage-Gated Sodium Channel Blockers |
| **Buflomedil** | Vasodilator Agents |
| **Levobunolol** | Adrenergic beta-Antagonists/Sympatholytics |
| **PHA-00851261E** | ? |
| **Estropipate** | Contraceptive Agents |
| **Ioversol** | Contrast Media |
| **0175029-0000** | ? |
| **Gelsemine** | Alkaloids/? |

**Table S3.** **List of 35 selected molecules.** CMAP concentration (µM) and calculated EC50 in the context of pre-treatment/treatment *in vitro*.

| Molecule | CMAP (μM) | EC50 (nM) |
| --- | --- | --- |
| Monensin | 6 | 3.27 |
| Biperiden | 11 | 0.38 |
| Felbinac | 19 | 34.35 |
| Sulfamonomethoxine | 14 | 0.28 |
| Adiphenine | 11 | 6.29 |
| Benzathin Benzylpenicilline | 4 | 0.26 |
| Gentamycin | 3 | 0.039 |
| Timolol | 9 | nd |
| Lanatoside C | 4 | nd |
| Alpha-estradiol | 0.01 | nd |
| Apigenin | 15 | 0.21 |
| Amiloride | 13 | 0.41 |
| Ribavirin | 16 | 0.86 |
| Methoxamine | 16 | nd |
| Tolazoline | 20 | 0.93 |
| Folic acid | 9 | 0.22 |
| Diphenhydramine | 14 | 0.25 |
| Roxythromycin | 5 | 0.14 |
| Pindolol | 16 | 0.79 |
| Prestwick-1103 | 20 | 626.63 |
| Carmustine | 100 | 0.07 |
| Clofilium tosylate | 8 | 0.16 |
| Prednisone | 11 | nd |
| Choropyramine | 12 | 0.07 |
| Urseodeoxycholic acid | 10 | 0.61 |
| Ranitidine | 11 | 0.68 |
| Flucytosine | 31 | 2.02 |
| Diltiazem | 9 | 0.84 |
| Fusidic acid | 7 | 0.038 |
| Levamisole | 17 | nd |
| Sulfadimethoxine | 13 | 0.69 |
| Riboflavine | 11 | nd |
| Genistein | 10 | 1.06 |
| Nimesulide | 13 | nd |
| Etilefrine | 18 | 0.34 |

**Table S4.**
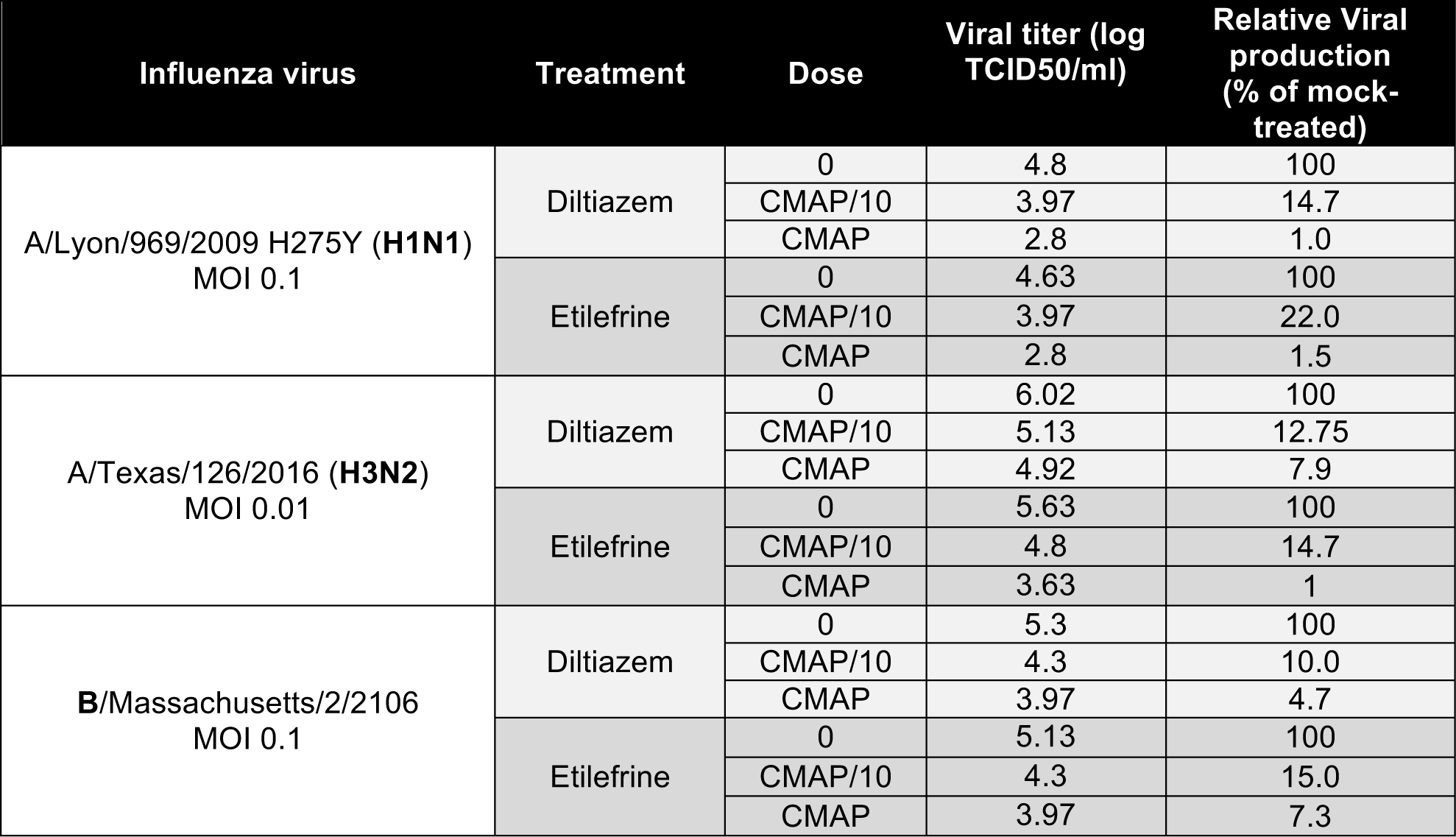
**Evaluation of antiviral efficacy of diltiazem or etilefrine in the context of infection by different influenza strains.** Human lung epithelial cells (A549) were incubated with supplemented medium (mock), or different concentrations of diltiazem (CMAP, 9 µM) or etilefrine (CMAP, 18 µM). Six hours after treatment, cells were washed and then infected with different prototype human influenza strains (as indicated). One hour after viral infection, a second identical treatment dose in supplemented medium was added. Relative viral titers compared to the mock-treated control are shown. Results are representative of two independent experiments, and confirm the antiviral activity of diltiazem and etilefrine on oseltamivir-resistant A(H1N1)pdm09, as well as wild-type H3N2 and B influenza strains.

**Table S5.** **Virus pre-incubation with diltiazem or etilefrine does not interfere with early viral entry steps.** Two viral dilutions (#1 and #2, respectively 10^6^ and 10^5^ TCID50/mL) were pre-incubated for 1 h with PBS, diltiazem (CMAP, 9 µM), etilefrine (CMAP, 18 µM), or oseltamivir (1 µM). A(H1N1)pdm09 positive and a negative sera were used as controls. After incubation, viral titers (log10 TCID50/mL) were determined in MDCK cells. Results are representative of two independent experiments and indicate that pre-incubation with either diltiazem or etilefrine does not affect viral titers compared to PBS-incubated control, suggesting that the antiviral effect of these molecules is not mediated by direct drug-virus interactions at early stages of viral entry.

| Pre-incubation treatment | Viral titer (log TCID<sub>50</sub>/ml) |  | Mean relative Viral production (% of mock-treated) |
| --- | --- | --- | --- |
|  | Dilution #1 | Dilution #2 |  |
| Control PBS | 5.3 | 4.63 | - |
| Diltiazem (9 μM) | 5.63 | 4.63 | 156.9 |
| Etilefrine (18 μM) | 5.30 | 4.63 | 100.0 |
| Oseltamivir (1 μM) | 5.63 | 4.30 | 130.3 |
| Negative serum | 4.97 | 4.30 | 47.7 |
| Positive serum | 3.30 | 2.53 | 0.9 |

**Table S6.** Apical viral production and transepithelial electrical resistance (TEER) in infected MucilAir^®^ human airway epithelium (HAE). MucilAir^®^ HAE were infected on the apical pole with influenza A/Lyon/969/09 (H1N1)pdm09 virus at a MOI of 0.1 and treated on the basolateral pole. Treatments were initiated 5 h after infection and were continued once daily for 4 additional days. *p<0.05, **p<0.01 and ***p<0.001 compared to the infected mock-treated group by one-way ANOVA with Tukey’s post-test. Data are representative of at least three independent experiments.

| Hours<br>P.I. | Treatment | Apical viral titer | Apical viral titer | $\Delta$ TEER |
| --- | --- | --- | --- | --- |
|  |  | PFU/ml | log TCID<sub>50</sub>/ml | Ohm/cm<sup>2</sup> |
|  |  | (CI95) | (CI95) | (CI95) |
| 24 | Mock | 7.2<sup>^</sup>5 (2.3<sup>^</sup>5 – 1.2<sup>^</sup>6) | 6.74 (6.53 – 6.95) | 59.84 (-35.75 – 155.4) |
| | Oseltamivir 0.1 μM | 2.1<sup>^</sup>5 (1.6<sup>^</sup>4 – 4.1<sup>^</sup>5) | 6.02 (4.69 – 7.35) | 8.98 (-81.72 – 99.68) |
| | Oseltamivir 1 μM | 2.1<sup>^</sup>4*** (2.3<sup>^</sup>3 – 3.9<sup>^</sup>4) | 5.13*** (4.61 – 5.66) | -24.32 (-58.00 – 9.36) |
| | Diltiazem 9 μM | 3.8<sup>^</sup>6 (1.6<sup>^</sup>5 – 6.0<sup>^</sup>5) | 6.47 (5.03 – 7.90) | 35.10 (10.96 – 59.23) |
| | Diltiazem 90 μM | 3.1<sup>^</sup>4*** (1.8<sup>^</sup>5 – 4.3<sup>^</sup>5) | 5.57** (4.95 – 6.18) | 86.13 (3.72 – 168.5) |
| | Ose 0.1 μM / Dil 9 μM | 5.3<sup>^</sup>4*** (1.7<sup>^</sup>3 – 1.0<sup>^</sup>5) | 6.25 (5.62 – 6.89) | 34.50 (-33.15 – 102.2) |
| | Ose 1 μM / Dil 90 μM | 2.3<sup>^</sup>4*** (1.9<sup>^</sup>4 – 2.7<sup>^</sup>4) | 5.19*** (4.15 – 6.23) | 32.96 (-77.24 – 143.2) |
| 48 | Mock | 2.1<sup>^</sup>8 (1.3<sup>^</sup>8 – 3.0<sup>^</sup>8) | 9.12 (8.91 – 9.33) | -2.64 (-233.3 – 228.0) |
| | Oseltamivir 0.1 μM | 5.9<sup>^</sup>7** (4.9<sup>^</sup>7 – 6.8<sup>^</sup>7) | 8.61 (7.43 – 9.80) | 20.09 (-39.75 – 79.94) |
| | Oseltamivir 1 μM | 8.8<sup>^</sup>4*** (3.2<sup>^</sup>3 – 1.7<sup>^</sup>5) | 6.20*** (5.39 – 7.01) | 7.26 (-85.73 – 100.3) |
| | Diltiazem 9 μM | 5.8<sup>^</sup>7** (3.0<sup>^</sup>7 – 8.5<sup>^</sup>7) | 8.91 (8.67 – 9.16) | -48.31 (-180.1 – 83.47) |
| | Diltiazem 90 μM | 1.0<sup>^</sup>5*** (4.4<sup>^</sup>4 – 1.5<sup>^</sup>5) | 6.56*** (5.23 – 7.88) | 120.1 (-85.77 – 326.00) |
| | Ose 0.1 μM / Dil 9 μM | 2.3<sup>^</sup>6*** (9.7<sup>^</sup>5 – 3.6<sup>^</sup>6) | 7.54* (6.92 – 8.17) | 78.28 (33.37 – 123.2) |
| | Ose 1 μM / Dil 90 μM | 2.6<sup>^</sup>4*** (1.4<sup>^</sup>4 – 3.8<sup>^</sup>4) | 5.38*** (5.05 – 5.71) | -8.84 (-58.73 – 41.05) |
| 72 | Mock | 1.2<sup>^</sup>8 (3.3<sup>^</sup>7 – 2.1<sup>^</sup>8) | 8.48 (8.02 – 8.94) | -244.1 (-275.2 – -213.0) |
| | Oseltamivir 0.1 μM | 4.6<sup>^</sup>7* (7.6<sup>^</sup>6 – 8.4<sup>^</sup>7) | 8.30 (7.73 – 8.87) | -110.9* (-256.0 – 34.2) |
| | Oseltamivir 1 μM | 1.7<sup>^</sup>5*** (7.1<sup>^</sup>4 – 2.7<sup>^</sup>5) | 6.85*** (6.14 – 7.56) | 22.55*** (-20.25 – 65.35) |
| | Diltiazem 9 μM | 1.9<sup>^</sup>7** (-4.7<sup>^</sup>5 – 3.9<sup>^</sup>7) | 8.21 (7.28 – 9.14) | -218.0 (255.6 – -180.4) |
| | Diltiazem 90 μM | 2.1<sup>^</sup>5*** (1.3<sup>^</sup>5 – 3.0<sup>^</sup>5) | 7.38* (6.53 – 8.22) | -115.5 (-308.5 – 77.52) |
| | Ose 0.1 μM / Dil 9 μM | 1.8<sup>^</sup>7** (6.9<sup>^</sup>6 – 2.8<sup>^</sup>7) | 8.41 (7.38 – 9.45) | -37.88** (-215.3 – 139.6) |
| | Ose 1 μM / Dil 90 μM | 4.7<sup>^</sup>4*** (3.8<sup>^</sup>4 – 5.7<sup>^</sup>4) | 5.62*** (5.23 – 6.01) | -43.16** (-77.17 – -9.15) |
| 96 | Mock | 2.8<sup>^</sup>7 (1.3<sup>^</sup>7 – 4.2<sup>^</sup>7) | 7.86 (7.74 – 7.99) | -267.3 (-288.7 – -246.0) |
| | Oseltamivir 0.1 μM | 4.1<sup>^</sup>6*** (-2.3<sup>^</sup>6 – 1.0<sup>^</sup>7) | 7.80 (6.56 – 9.04) | -209.2 (-271.5 – -146.8) |
| | Oseltamivir 1 μM | 1.8<sup>^</sup>5*** (6.3<sup>^</sup>3 – 3.4<sup>^</sup>5) | 7.09 (5.79 – 8.40) | 10.67*** (-27.47 – 48.81) |
| | Diltiazem 9 μM | 2.3<sup>^</sup>6*** (1.9<sup>^</sup>6 – 2.7<sup>^</sup>6) | 7.32 (6.43 – 8.22) | -220.5 (-267.4 – -173.5) |
| | Diltiazem 90 μM | 1.6<sup>^</sup>5*** (-5.6<sup>^</sup>4 – 3.7<sup>^</sup>5) | 7.37 (5.97 – 8.76) | 24.59*** (-10.79 – 59.97) |
| | Ose 0.1 μM / Dil 9 μM | 4.0<sup>^</sup>6*** (1.2<sup>^</sup>6 – 6.8<sup>^</sup>6) | 7.84 (7.38 – 8.31) | -149.9* (-379.8 – 80.1) |
| | Ose 1 μM / Dil 90 μM | 2.9<sup>^</sup>4*** (2.1<sup>^</sup>4 – 3.7<sup>^</sup>4) | 5.47** (4.38 – 6.56) | -59.79*** (-97.62 – -21.96) |

## References

1. Influenza facts Sheet n°211. 2014. http://www.who.int/mediacentre/factsheets/fs211/en/. Accessed 20 Jan 2016.

2. Loregian A, Mercorelli B, Nannetti G, Compagnin C, Palù G. ntiviral strategies against influenza virus: towards new therapeutic approaches. Cell Mol Life Sci CMLS. 2014;71:3659–83.

3. Booth B, Zemmel R. Prospects for productivity. Nat Rev Drug Discov. 2004;3:451–6.

4. Ashburn TT, Thor KB. Drug repositioning: identifying and developing new uses for existing drugs. Nat Rev Drug Discov. 2004;3:673–83.

5. Law GL, Tisoncik-Go J, Korth MJ, Katze MG. Drug repurposing: a better approach for infectious disease drug discovery? Curr Opin Immunol. 2013;25:588–92.

6. Johansen LM, DeWald LE, Shoemaker CJ, Hoffstrom BG, Lear-Rooney CM, Stossel A, et al. A screen of approved drugs and molecular probes identifies therapeutics with anti-Ebola virus activity. Sci Transl Med. 2015;7:290ra89.

7. Xu M, Lee EM, Wen Z, Cheng Y, Huang W-K, Qian X, et al. Identification of small-molecule inhibitors of Zika virus infection and induced neural cell death via a drug repurposing screen. Nat Med. 2016;22:1101–7.

8. Naylor S, Schonfeld JM. Therapeutic drug repurposing, repositioning and rescue - Part I: Overview. Drug Discov World. 2014;16. https://www.researchgate.net/publication/286670692_Therapeutic_drug_repurposing_repositioning_and_rescue_-_Part_I_Overview. Accessed 11 Jan 2017.

9. Keiser MJ, Setola V, Irwin JJ, Laggner C, Abbas AI, Hufeisen SJ, et al. Predicting new molecular targets for known drugs. Nature. 2009;462:175–81.

10. Haupt VJ, Schroeder M. Old friends in new guise: repositioning of known drugs with structural bioinformatics. Brief Bioinform. 2011;12:312–26.

11. Li Y, Agarwal P. A pathway-based view of human diseases and disease relationships. PloS One. 2009;4:e4346.

12. Lussier YA, Chen JL. The emergence of genome-based drug repositioning. Sci Transl Med. 2011;3:96ps35.

13. Hütter G, Bodor J, Ledger S, Boyd M, Millington M, Tsie M, et al. CCR5 Targeted Cell Therapy for HIV and Prevention of Viral Escape. Viruses. 2015;7:4186–203.

14. Ludwig S. Disruption of virus-host cell interactions and cell signaling pathways as an anti-viral approach against influenza virus infections. Biol Chem. 2011;392:837–47.

15. Rossignol JF, La Frazia S, Chiappa L, Ciucci A, Santoro MG. Thiazolides, a new class of anti-influenza molecules targeting viral hemagglutinin at the post-translational level. J Biol Chem. 2009;284:29798–808.

16. Belser JA, Lu X, Szretter KJ, Jin X, Aschenbrenner LM, Lee A, et al. DAS181, a novel sialidase fusion protein, protects mice from lethal avian influenza H5N1 virus infection. J Infect Dis. 2007;196:1493–9.

17. Mazur I, Wurzer WJ, Ehrhardt C, Pleschka S, Puthavathana P, Silberzahn T, et al. Acetylsalicylic acid (ASA) blocks influenza virus propagation via its NF-kappaB-inhibiting activity. Cell Microbiol. 2007;9:1683–94.

18. Josset L, Textoris J, Loriod B, Ferraris O, Moules V, Lina B, et al. Gene expression signature-based screening identifies new broadly effective influenza a antivirals. PloS One. 2010;5.

19. Lamb J, Crawford ED, Peck D, Modell JW, Blat IC, Wrobel MJ, et al. The Connectivity Map: using gene-expression signatures to connect small molecules, genes, and disease. Science. 2006;313:1929–35.

20. Lamb J. The Connectivity Map: a new tool for biomedical research. Nat Rev Cancer. 2007;7:54–60.

21. Escuret V, Cornu C, Boutitie F, Enouf V, Mosnier A, Bouscambert-Duchamp M, et al. Oseltamivir-zanamivir bitherapy compared to oseltamivir monotherapy in the treatment of pandemic 2009 influenza A(H1N1) virus infections. Antiviral Res. 2012;96:130–7.

22. Khaznadar Z, Boissel N, Agaugué S, Henry G, Cheok M, Vignon M, et al. Defective NK Cells in Acute Myeloid Leukemia Patients at Diagnosis Are Associated with Blast Transcriptional Signatures of Immune Evasion. J Immunol Baltim Md 1950. 2015;195:2580–90.

23. Degletagne C, Keime C, Rey B, de Dinechin M, Forcheron F, Chuchana P, et al. Transcriptome analysis in non-model species: a new method for the analysis of heterologous hybridization on microarrays. BMC Genomics. 2010;11:344.

24. Tukey J, Wilder J. Exploratory Data Analysis. Addison-Wesley; 1977.

25. Benjamini Y, Hochberg Y. Controlling the False Discovery Rate: A Practical and Powerful Approach to Multiple Testing. J R Stat Soc Ser B Methodol. 1995;57:289–300.

26. Barrett T, Edgar R. Gene expression omnibus: microarray data storage, submission, retrieval, and analysis. Methods Enzymol. 2006;411:352–69.

27. Dennis G Jr, Sherman BT, Hosack DA, Yang J, Gao W, Lane HC, et al. DAVID: Database for Annotation, Visualization, and Integrated Discovery. Genome Biol. 2003;4:P3.

28. Ashburner M, Ball CA, Blake JA, Botstein D, Butler H, Cherry JM, et al. Gene ontology: tool for the unification of biology. The Gene Ontology Consortium. Nat Genet. 2000;25:25–9.

29. Hatakeyama S, Sakai-Tagawa Y, Kiso M, Goto H, Kawakami C, Mitamura K, et al. Enhanced expression of an alpha2,6-linked sialic acid on MDCK cells improves isolation of human influenza viruses and evaluation of their sensitivity to a neuraminidase inhibitor. J Clin Microbiol. 2005;43:4139–46.

30. Moules V, Ferraris O, Terrier O, Giudice E, Yver M, Rolland JP, et al. In vitro characterization of naturally occurring influenza H3NA-viruses lacking the NA gene segment: toward a new mechanism of viral resistance? Virology. 2010;404:215–24.

31. Tsai AW, McNeil CF, Leeman JR, Bennett HB, Nti-Addae K, Huang C, et al. Novel Ranking System for Identifying Efficacious Anti-Influenza Virus PB2 Inhibitors. Antimicrob Agents Chemother. 2015;59:6007–16.

32. Bolger AM, Lohse M, Usadel B. Trimmomatic: a flexible trimmer for Illumina sequence data. Bioinforma Oxf Engl. 2014;30:2114–20.

33. Bray NL, Pimentel H, Melsted P, Pachter L. Near-optimal probabilistic RNA-seq quantification. Nat Biotechnol. 2016;34:525–7.

34. Robinson MD, McCarthy DJ, Smyth GK. edgeR: a Bioconductor package for differential expression analysis of digital gene expression data. Bioinforma Oxf Engl. 2010;26:139–40.

35. Dupinay T, Nguyen A, Croze S, Barbet F, Rey C, Mavingui P, et al. Next-generation sequencing of ultra-low copy samples: From clinical FFPE samples to single-cell sequencing. Curr Top Virol. 2013;10:63–83.

36. Kinsella RJ, Kähäri A, Haider S, Zamora J, Proctor G, Spudich G, et al. Ensembl BioMarts: a hub for data retrieval across taxonomic space. Database J Biol Databases Curation. 2011;2011:bar030.

37. Flicek P, Amode MR, Barrell D, Beal K, Billis K, Brent S, et al. Ensembl 2014. Nucleic Acids Res. 2014;42 Database issue:D749–755.

38. Durr FE, Lindh HF. Efficacy of ribavirin against influenza virus in tissue culture and in mice. Ann N Y Acad Sci. 1975;255:366–71.

39. Leyssen P, De Clercq E, Neyts J. Molecular strategies to inhibit the replication of RNA viruses. Antiviral Res. 2008;78:9–25.

40. Alonso FV, Compans RW. Differential effect of monensin on enveloped viruses that form at distinct plasma membrane domains. J Cell Biol. 1981;89:700–5.

41. Hayden F. Developing new antiviral agents for influenza treatment: what does the future hold? Clin Infect Dis Off Publ Infect Dis Soc Am. 2009;48 Suppl 1:S3–13.

42. Lee SM-Y, Yen H-L. Targeting the host or the virus: current and novel concepts for antiviral approaches against influenza virus infection. Antiviral Res. 2012;96:391–404.

43. Koh Y. Update in acute respiratory distress syndrome. J Intensive Care. 2014;2:2.

44. Poissy J, Terrier O, Lina B, Textoris J, Rosa-Calatrava M. La modulation de la signature transcriptomique de l’hôte infecté?: une nouvelle stratégie thérapeutique dans les viroses graves?? Exemple de la grippe. Réanimation. 2016;25:53–61.

45. Lou Z, Sun Y, Rao Z. Current progress in antiviral strategies. Trends Pharmacol Sci. 2014;35:86–102.

46. Dayekh K, Johnson-Obaseki S, Corsten M, Villeneuve PJ, Sekhon HS, Weberpals JI, et al. Monensin inhibits epidermal growth factor receptor trafficking and activation: synergistic cytotoxicity in combination with EGFR inhibitors. Mol Cancer Ther. 2014;13:2559–71.

47. Zhang Y, Jamaluddin M, Wang S, Tian B, Garofalo RP, Casola A, et al. Ribavirin treatment up-regulates antiviral gene expression via the interferon-stimulated response element in respiratory syncytial virus-infected epithelial cells. J Virol. 2003;77:5933–47.

48. Pubchem C10H15NO2. Etilefrine | C10H15NO2 - PubChem. https://pubchem.ncbi.nlm.nih.gov/compound/Etilefrine. Accessed 3 Feb 2017.

49. Pubchem C22H26N2O4S. diltiazem | C22H26N2O4S - PubChem. https://pubchem.ncbi.nlm.nih.gov/compound/diltiazem. Accessed 3 Feb 2017.

50. Fujioka Y, Nishide S, Ose T, Suzuki T, Kato I, Fukuhara H, et al. A Sialylated Voltage-Dependent Ca2+ Channel Binds Hemagglutinin and Mediates Influenza A Virus Entry into Mammalian Cells. Cell Host Microbe. 2018;23:809–818.e5.

